# High-throughput generation and phenotypic characterization of zebrafish CRISPR mutants of DNA repair genes

**DOI:** 10.1101/2020.10.04.325621

**Authors:** Unbeom Shin, Khriezhanuo Nakhro, Chang-Kyu Oh, Blake Carrington, Hayne Song, Gaurav Varshney, Youngjae Kim, Hyemin Song, Sangeun Jeon, Gabrielle Robbins, Sangin Kim, Suhyeon Yoon, Yongjun Choi, Suhyung Park, Yoo Jung Kim, Shawn Burgess, Sukhyun Kang, Raman Sood, Yoonsung Lee, Kyungjae Myung

## Abstract

A systematic knowledge of the roles of DNA repair genes at the level of the organism has been limited due to the lack of appropriate experimental techniques. Here, we generated zebrafish loss-of-function mutants for 32 DNA repair and replication genes through multiplexed CRISPR/Cas9-mediated mutagenesis. High-throughput phenotypic characterization of our mutant collection revealed that three genes (*atad5a*, *ddb1, pcna*) are essential for proper embryonic development and hematopoiesis; seven genes (*apex1*, *atrip*, *ino80*, *mre11a*, *shfm1*, *telo2*, *wrn*) are required for growth and development during juvenile stage and six genes (*blm*, *brca2*, *fanci, rad51*, *rad54l*, *rtel1*) play critical roles in sex development. Furthermore, mutation in six genes (*atad5a*, *brca2*, *polk*, *rad51*, *shfm1*, *xrcc1*) displayed hypersensitivity to DNA damage agents. Further characterization of *atad5a*^*−/−*^ mutants demonstrate that Atad5a is required for normal brain development and hematopoiesis. Our zebrafish mutant collection provides a unique resource for understanding of the roles of DNA repair genes at the organismal level.

## INTRODUCTION

DNA repair is critical for the maintenance of genomic integrity in living organisms. Our genomes continuously encounter various kinds of challenges such as oxidation, deamination, alkylation, and formation of bulky adducts, mismatches and DNA breaks. Those damages are derived from endogenous sources such as reactive oxygen species, alkylating agents and replication errors, and exogenous agents including ionizing radiation (IR), ultraviolet radiation and environmental chemicals (Chatterjee and Walker, 2017). To protect our genome from those challenges, cellular systems employ several DNA repair pathways including base excision repair (BER), nucleotide excision repair (NER), mismatch repair (MMR), translesion synthesis (TLS), homologous recombination (HR) and non-homologous end-joining (NHEJ) (Iyama and Wilson, 2013, Friedberg, 2003). These mechanisms have been widely studied at molecular and cellular level, but a systematic investigation at the organismal level is not currently available.

Loss-of-function studies using knock-out (KO) mice have revealed numerous functional roles of DNA repair processes during vertebrate development (Specks et al., 2015). For example, normal neurogenesis requires distinct DNA damage and repair processes. Genetic defects in ATM, which activates DNA double-strand break (DSB) damage responses, cause abnormalities in the nerve system in adult mice (Shiloh and Ziv, 2013, Barlow et al., 1996) and mutation of DNA-PKcs required for NHEJ leads to aberrant neurogenesis affecting neural progenitor cells and post-mitotic neurons (Enriquez-Rios et al., 2017). Moreover, certain DNA repair processes are involved in distinct mice developmental processes. Fanconi anemia (FA) genes critical for DNA interstrand cross-link (ICL) repair showed their diverse roles during embryogenesis. KO mice of *Fanca* and *Fancc* showed abnormal development of hematopoietic progenitors, while *Fanca* and *Fancg* mutant mice demonstrated microcephaly with induced apoptosis of embryonic neural progenitors (Rathbun et al., 1997, Rio et al., 2002, Sii-Felice et al., 2008). These FA gene KO mice also showed defects of germ cell development with hypogonadism (Parmar et al., 2009). In addition, conditional KO studies revealed the function of genes related to DNA damage and repair processes on specific tissue or organ development. Mice deficient in the ubiquitin ligase adapter Ddb1 (DNA damage binding protein 1) in the brain causes abnormal brain growth and hemorrhages, while blood lineage-specific deletion of *Ddb1* leads to failure of hematopoietic stem cell development (Cang et al., 2006, Gao et al., 2015).

Although numerous KO mice of genes involved in DNA repair have been generated, uncovering how these genes affect developmental processes is hampered by the early embryonic lethality associated with mouse KO mutations in a large number of DNA repair genes (Suzuki et al., 1997, Lim and Hasty, 1996, Cang et al., 2006, Buis et al., 2008, Qiu et al., 2016, Ding et al., 2004). For further understanding of gene function, tissue-specific or inducible genetic mutants in mice are critical for the study of DNA repair and replication (Cang et al., 2006, Lee et al., 2009). Teleost zebrafish has emerged as a powerful alternative genetic model system to mice, possessing similar DNA repair systems as mammals (Pei and Strauss, 2013). A key advantage of zebrafish is their faster development bypassing possible early embryonic lethality of essential genes in zygotic KO mutants due to maternal mRNA deposition effects by early gastrulation stage, compared to mice system where maternal mRNA is mostly degraded before 2-cell stage (Svoboda and Flemr, 2010, Tadros and Lipshitz, 2009). Zebrafish therefore provide an optimal model to investigate the roles of genes critical during early embryogenesis. Moreover, recently reported multiplexed mutagenesis using CRISPR/Cas9 system allows for the generation of targeted gene mutation on a large scale (Varshney et al., 2015). Finally, mutant phenotypes can be continuously examined throughout development in a high-throughput manner due to the visual transparency of embryos during embryogenesis and high fecundity from single zebrafish breeding (Fuentes et al., 2018).

Here, we generated 65 mutant alleles disrupting 32 genes involved in DNA repair and replication using multiplexed CRISPR/Cas9 mutagenesis and characterized their phenotypes in embryonic development, viability, growth, germ cell development, hematopoiesis and sensitivity to DNA damage. Our data revealed that deficiency of 18 of these 32 DNA repair genes lead to at least one phenotype in the homozygous mutant animals, thus revealing their critical roles in embryogenesis, hematopoiesis, growth during larval and juvenile stages, germ cell development and response to DNA damaging agents. We chose *atad5a* mutants, deficient in unloading the DNA clamp PCNA upon completion of DNA replication and repair, for more detailed phenotypic analysis and determined how *atad5a* deficiency leads to reduced brain size and hematopoietic defects in *atad5a* mutants. Our data showed that reduced brain structure and diminished number of hematopoietic stem and progenitor cells (HSPCs) are mainly caused by p53-induced apoptosis in the brain and caudal hematopoietic tissue (CHT), respectively and they can be rescued by inhibition of apoptosis by *p53* ablation or inhibition of ATM signaling. We therefore directly show that ATM-p53-dependent apoptosis is induced in the brain and blood population during embryogenesis without normal function of Atad5a and possible other crucial DNA repair genes.

## Results

### Multiplexed CRISPR mutagenesis generated 32 loss-of-function zebrafish mutants of DNA repair genes

Aiming to mutate 56 genes related to DNA repair and replication processes, multiplexed injections of sgRNAs with *Cas9* mRNA in 16 injection groups (Fig. 1A; Table S1) were performed as previously described (Varshney et al., 2015). Up to four gRNAs targeting distinct gene loci from different chromosomes were injected into fertilized zebrafish eggs at 1-cell stage with *Cas9* mRNA in each injection group. To identify whether potential in/del mutations generated by CRIPSR/Cas9 system are germline-transmitted, F1 progeny embryos from injected candidate founder fish were analyzed through genetic fragment analysis with fluorescence PCR for each gene in the injection group. We first identified in/del mutations in 49 genes, indicating that sgRNAs to 7 genes failed to induce mutations. Next, individual F1 animals carrying mutations were crossed with wild-type to generate F2 germline-transmitted mutant families, retrieving mutations in 34 genes, while mutant candidates with the other 15 genes failed to inherit potential in/del mutations. The siblings of F2 heterozygous mutants were then crossed to produce homozygous mutants for each allele, and genotyped again. Eventually, we recovered mutant animals containing 67 mutant alleles for 34 genes involved in DNA repair processes and replication (Table S2). Among 34 genes, mutants of 2 genes (*polh* and *palb2*) only possessed in/del alleles with in-frame mutations, while other 32 genes were successfully mutated, generating multiple alleles with frameshift mutations that cause premature stop codons (Table 1).

**Fig. 1.**
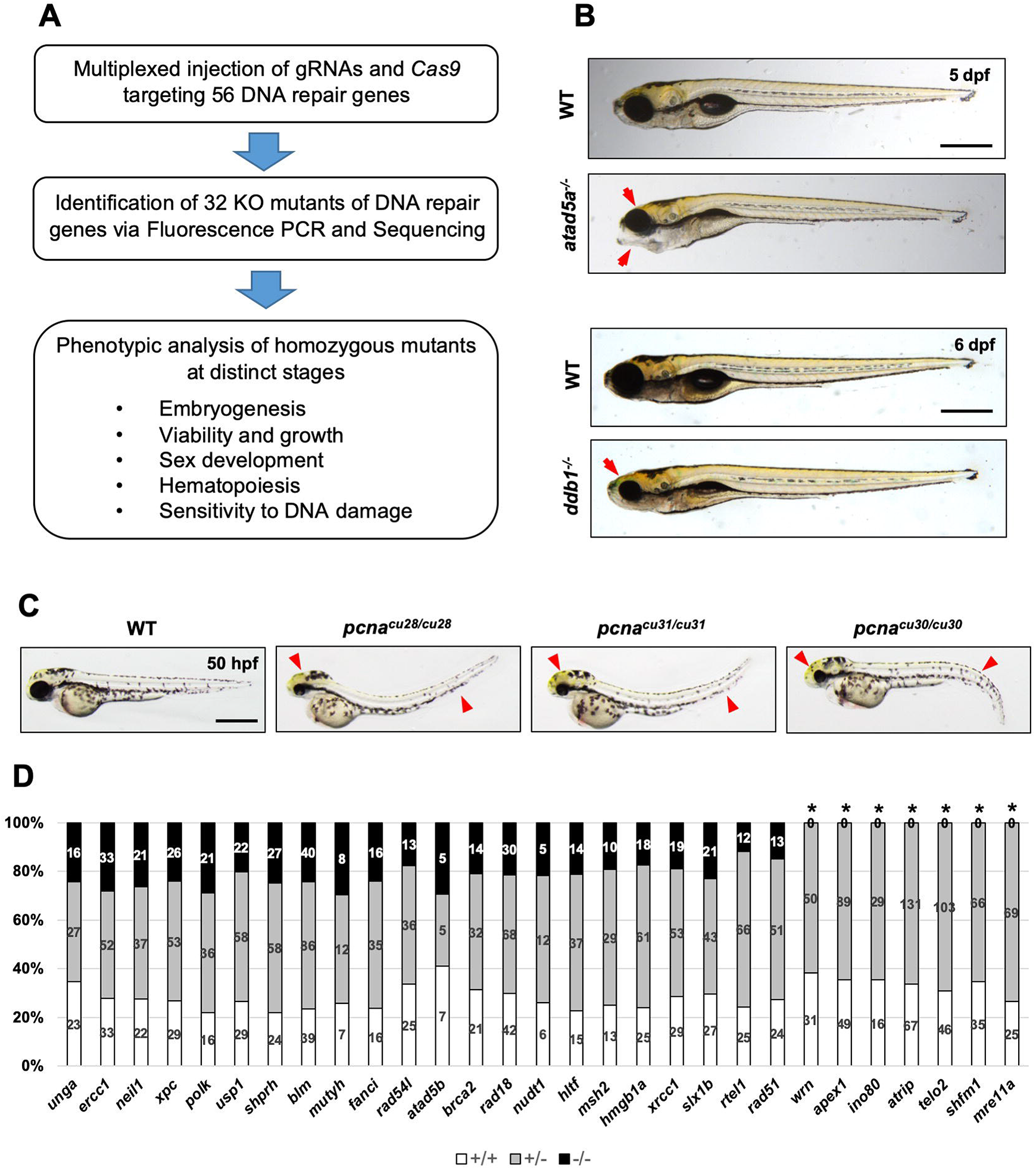
Large-scale phenotypic characterization of zebrafish CRISPR mutants related to DNA repair genes. **A.** Overview of experimental procedures for generation and phenotypic characterization of the zebrafish mutants of DNA repair genes. **B.** Lateral view of mutant embryos of *atad5a*^*−/−*^(*cu33/cu33*) at 5 dpf and *ddb1*^*−/−*^(*cu43/cu43*) at 6 dpf with their sibling WT controls. Red arrowheads indicate morphological alterations in the mutant embryos. **C.** Images of *pcna* mutants and WT control at 50 hpf. *pcna*^*cu28/cu28*^ possessed 1 base pair (bp) deletion producing null mutants, while *pcna*^*cu30/cu30*^ and *pcna*^*cu31/cu31*^ contain in-frame deletion generating deleted protein of PCNA monomer of Δ^46^SL^47^ and Δ^44^HVS^46^, respectively. Red arrowheads indicate morphological alterations in the mutant embryos. **D.** A graph describes percentage of fish progeny with genotype (+/+, +/−, −/−) from each inbred heterozygous mutant line at 4 to 6 months of age. Numbers in each graph bar represent the number of fish with each genotype. Black asterisks indicate the mutant alleles in which homozygous mutants are lethal at adult stage. Scale bar = 500 μm.

**Table 1.**
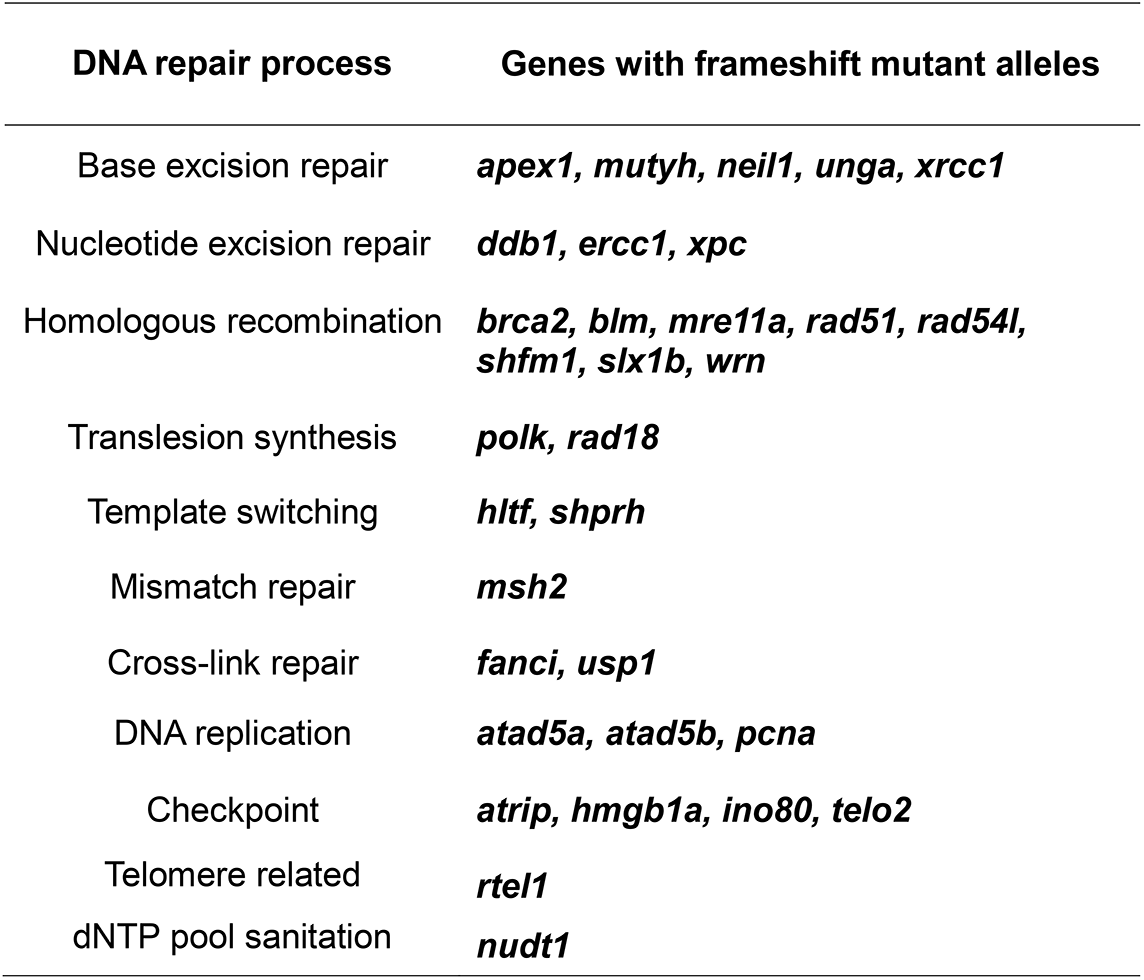
List of genes related to DNA repair and replication with frameshift mutations obtained from CRISPR mutagenesis.

### Phenotypic evaluation of the mutants of 32 DNA repair genes

Previously, deletion of a number of DNA repair genes in mice KO system resulted in lethality in early embryogenesis (e.g. *Apex1*, *Atad5*, *Blm*, *Brca2*, *Ddb1*, *Ino80*, *Mre11a*, *Rad51*, *Rtel1*, *Telo2*, *Xrcc1*) (Cang et al., 2006, Takai et al., 2007, Qiu et al., 2016, Ding et al., 2004, Buis et al., 2008, Lim and Hasty, 1996, Chester et al., 1998, Bell et al., 2011, Ludwig et al., 1998, Yan et al., 2004, Tebbs et al., 1999). To confirm its critical functions during early development and further characterize the roles of DNA repair genes spatiotemporally, we screened phenotypic alterations of homozygous mutants generated from our multiplexed CRISPR mutagenesis in distinct categories, including 1) embryonic development, 2) adult survival and growth, 3) sex determination and germ cell development, 4) hematopoiesis and 5) sensitivity to genotoxic stress (Fig. 1A). For phenotype screening, we utilized mutants of 32 genes with 51 loss-of-function alleles causing frameshift mutations (Table S3). All the mutants with defective phenotypes were only the ones with frameshift mutations except *pcna*^*−/−*^ as described below.

### Embryogenesis

Homozygous mutant embryos for three out of 32 genes showed morphological alterations followed by death before 7 dpf. *atad5a*^*cu33/cu33*^ mutants showed aberrant head development with reduced size of the brain at 5 dpf, while *ddb1*^*cu43/cu43*^ started to display anterior malformation including abnormalities of eye and jaw structure from 6 dpf compared to wild-type (WT) (Fig. 1B). Not surprisingly, in the absence of normal PCNA function, abnormal morphological appearance with reduced size of head and eyes as well as curved body trunk formation was observed at the earliest 50 hpf among all the mutant lines (Fig. 1C). Interestingly, the *pcna*^*cu30/cu30*^ and *pcna*^*cu31/cu31*^ mutants containing in-frame in/del mutations also displayed abnormal development (Fig. 1C; Fig. S1A). Immunoprecipitation analysis of the Pcna^cu31*/*cu31^ (Δ ^44^HVS^46^ mutation) cells showed that the protein was unstable to form the functional PCNA complex in those cells, while the Pcna^cu30*/*cu30^ (Δ ^46^SL^47^ mutation) failed to form homo-trimer essential for clamp formation in DNA replication (Fig. S1B and C). Due to lack of functional protein, *pcna*^*cu31/cu31*^ replicates defective phenotypes of null *pcna*^*cu28/cu28*^, while *pcna*^*cu30/cu30*^ showed distinctive morphology in the posterior part of the body, potentially suggesting multiple roles of PCNA during embryogenesis.

### Survival and growth

In contrast to what was observed in mice, our embryonic screening demonstrated that zebrafish mutants of majority of the DNA repair and replication genes develop normally during early development. Morphological defects were only obvious at later than 2 dpf, when the basic development is already completed (Fig. 1). To further characterize potential phenotypic defects, we examined viability of homozygous mutant animals at 4-6 months of age when the adult zebrafish become fully mature to produce progeny. Excluding the three mutants with embryonic defects introduced above, progenies generated from inbred heterozygous parents for 29 genes were genotyped to determine whether the homozygous mutant animals survived at expected Mendelian ratio. We found that homozygous mutants in seven genes: *apex1*^*−/−*^, *atrip*^*−/−*^, *shfm1*^*−/−*^, *ino80*^*−/−*^, *mre11a*^*−/−*^, *telo2*^*−/−*^, and *wrn*^*−/−*^ failed to survive to adult stages (Fig. 1D). To closely understand phenotypes of these mutants, we examined a lethal time-point for each mutant by identifying the developmental stage at which the number of homozygous mutants was significantly reduced. Interestingly, each mutant showed reduced survival rate at distinctive developmental stage. Homozygous mutants of *telo2*^*cu41/cu41*^ and *shfm1*^*cu46/cu46*^ died by 2 weeks post fertilization (Fig. S2). We further monitored possible morphological alterations and found that *telo2*^*cu41/cu41*^ embryos showed malformation of the head with a retrograded branchial arch as well as heart structure (Fig. 2A). Similarly, *shfm1*^*cu46/cu46*^ displayed defective head development with lens malformation. *apex1*^*cu2/cu2*^ and *atrip*^*cu37/cu37*^ mutants were not viable at 40 dpf, *mre11a*^*cu56/cu56*^ and *wrn*^*cu64/cu64*^ failed to survive to 60 dpf, while *ino80*^*cu36/cu36*^ scarcely survived to 90 dpf (Fig. S2). These five mutants displayed consistent phenotypes with reduced body size compared to their WT siblings at each designated developmental stage close to lethal time-point, suggesting potential function of each gene on either growth or physiology in vertebrates (Fig. 2B).

**Fig. 2.**
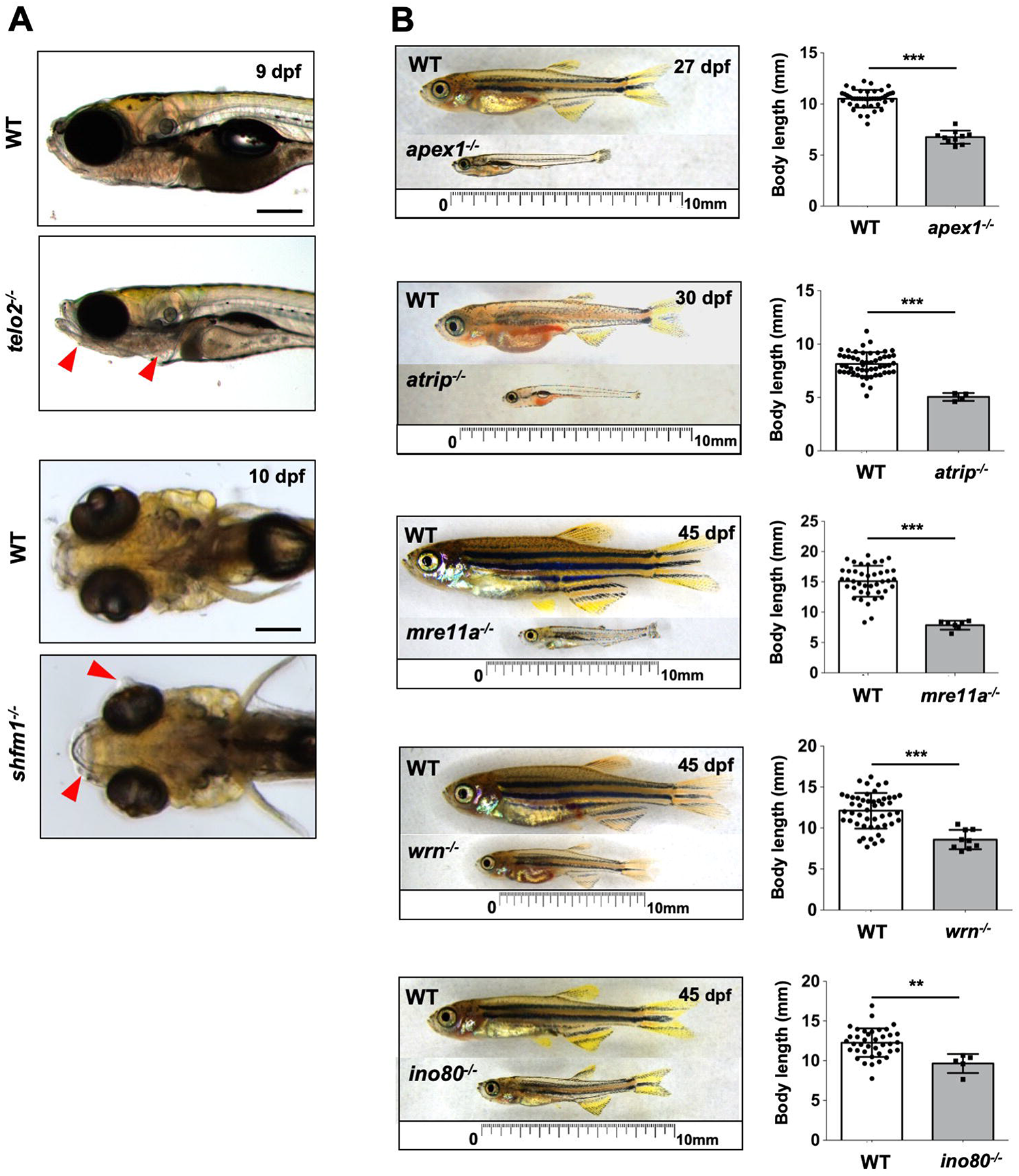
Screening of viability and morphology of mutants during larval and juvenile stages. **A.** Images of anterior region in the *telo2*^*−/−*^(*cu41/cu41*) and *shfm1*^*−/−*^(*cu46/cu46*) mutants and their WT sibling controls. Lateral view of *telo2*^*−/−*^(*cu41/cu41*) at 9 dpf showed abnormal formation of jawbone and heart structure, while dorsal view of *shfm1*^*−/−*^ (*cu46/cu46*) at 10 dpf demonstrated malformation of head part including mandible and lens. Red arrowheads indicate aberrant morphological structures in the mutants. **B.** Representative phenotypic images of each mutant (*apex1*^*−/−*^(*cu2/cu2*), *atrip*^*−/−*^(*cu37/cu37*), *mre11a*^*−/−*^(*cu56/cu56*), *wrn*^*−/−*^(*cu64/cu64*) and *ino80*^*−/−*^(*cu36/cu36*)) and its WT sibling controls at the designated time and quantification of the body length (fork length) of each mutant and WT control. Scale bars in each image indicate 10 mm. Graphs represent mean ± S.E.M. with individual values. p-values were calculated by unpaired two-tailed Student’s t-test. *** p<0.001, **p<0.01. Black scale bars = 200 μm.

### Sex reversal and germ cell development

Previous studies demonstrated that zebrafish mutants of HR and FA genes failed to generate female animals due to sex reversal defects without normal germ cell development (Shive et al., 2010, Botthof et al., 2017, Ramanagoudr-Bhojappa et al., 2018). To examine whether the DNA repair genes investigated here are involved in sex determination and germ cell development, we screened sexes of the 21 homozygous mutants that survive to 4-6 month of age (Fig. S3A). We found that all or most adult homozygous mutants were males in zebrafish lacking *blm, brca2*, *fanci, rad51*, *rad54l*, and *rtel1*, suggesting that sex reversal phenotypes developed in the absence of normal function of these genes (Fig. 3A; Fig. S3B). Further fertility tests for these six mutants showed that male homozygotes of *brca2*^*−/−*^, *rad51*^*−/−*^ and *blm*^*−/−*^ were infertile (fertilization success rate <6%), while the *rtel1*^*−/−*^, *fanci*^*−/−*^ and *rad54l*^*−/−*^ male animals were found to be fertile (fertilization success rate >65%) based on outcrosses with WT female animals (Fig. 3B; Table S4). Consistent with the finding of aberrant spermatogenesis due to meiotic arrest in *brca2*^*Q658X/Q658X*^ males (Shive et al., 2010), testes of our *brca2*^*cu51/cu51*^ males were void of spermatozoa, whereas WT siblings had normal spermatogenesis (Fig. 3C). Likewise, *blm*^*cu53/cu53*^ mutants exhibited the same skewed ratio with all homozygotes being males with possible sex reversal. Furthermore, although testis histology of *blm*^*cu53/cu53*^ homozygotes showed spermatogonia and an abundance of spermatocytes, the fraction of spermatozoa was severely diminished, indicating a failure of complete spermatogenesis similar to what was observed in *brca2*^*cu51/cu51*^ homozygotes (Fig. 3C).

**Fig. 3.**
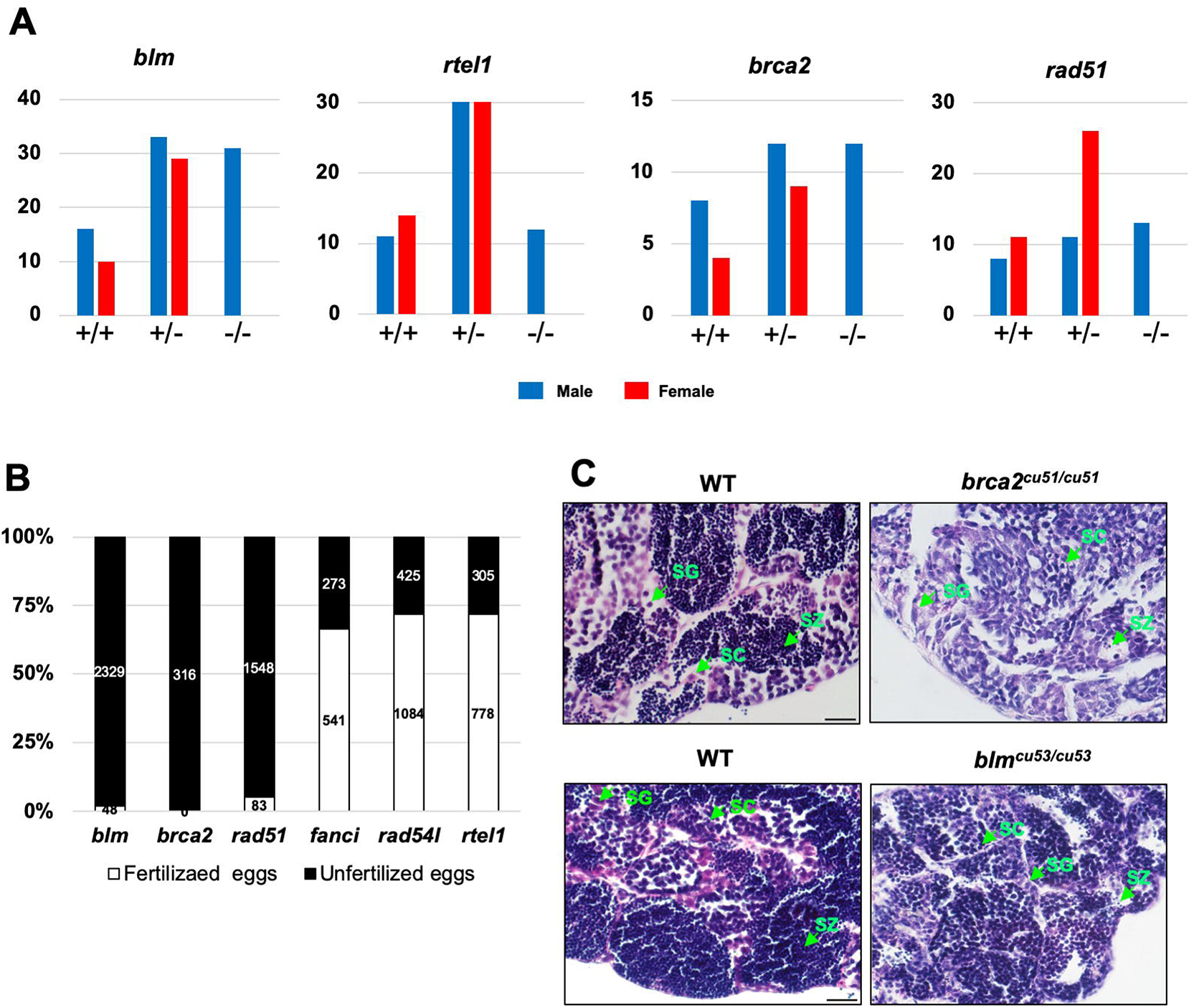
Mutants with sex reversal and abnormal germ cell development. **A**. The number of male and female animals genotyped from inbred heterozygous mutants of *blm, rtel1, brca2 and rad51*. **B**. Fertilization success rate of progenies generated from crosses between male homozygous mutants (*blm*^*−/−*^, *brca2*^*−/−*^, *rad51*^*−/−*^, *fanci*^*−/−*^, *rad54l*^*−/−*^ and *rtel1*^*−/−*^) and wild-type female zebrafish. **C.** Hematoxylin and eosin (H&E) staining of testis sections from the mutants of *brca2*^*cu51/cu51*^ and *blm*^*cu53/cu53*^, and their WT sibling controls. Green arrows indicate spermatogonia (sg), spermatocytes (sc) and mature spermatozoa (sz). Scale bar = 20 μm.

### Hematopoiesis

Although several studies have suggested that many DNA repair and replication genes play a key role in hematopoiesis and human blood disorders (Botthof et al., 2017, Gao et al., 2015, Alvarez et al., 2015, Xu et al., 2006, Marsh et al., 2018), their functions in hematopoietic progenitor development is not well understood. To understand how HSPC development is affected in our DNA repair mutant library, we examined expression of HSPC marker *cmyb* by whole-mount in situ hybridization (WISH). Hypothesizing that normal function of HSPCs is critical for animal survival, we assessed HSPC marker expression of 18 homozygous mutant embryos that show developmental and viability defects (Fig. 4A; Fig. S4A). WISH results showed that the *cmyb*-positive HSPC population in the CHT where HSPCs expand to be functional at 3-5 dpf (Tamplin et al., 2015) was dramatically reduced in three mutants that showed morphological defects before 6 dpf; *ddb1*^*cu43/cu43*^, *atad5a*^*cu33/cu33*^ and *pcna*^*cu29/cu29*^ (Fig. 4A). *cmyb* expression in the dorsal aorta where HSPCs emerge at 36 hpf was normal in these mutants, suggesting maintenance and expansion of HSPCs were affected, while HSPC emergence and specification was normal (Fig. S4B). We assessed proliferation and apoptosis through EdU-incorporation assays and acridine orange (AO) staining (Tucker and Lardelli, 2007), respectively, in these animals. In the *ddb1*^*cu43/cu43*^ mutants, proliferation was significantly reduced in the CHT without alteration of cell death events (Fig. 4B and C). In contrast, both *pcna*^*cu29/cu29*^ and *atad5a*^*cu33/cu33*^ mutants showed increased apoptosis as well as perturbed proliferation in the CHT, suggesting that they may affect HSPC development in a distinct manner.

**Fig. 4.**
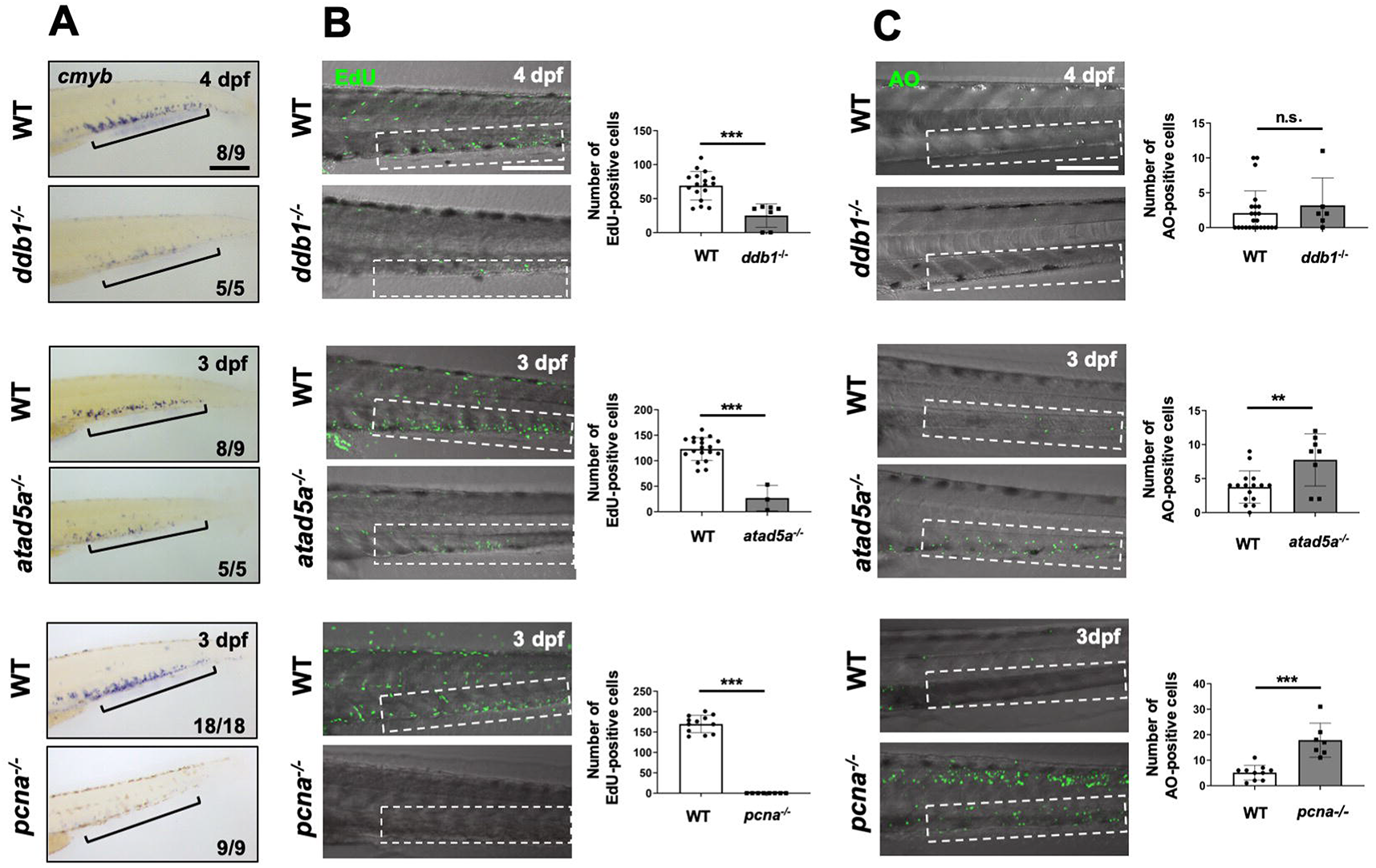
HSPC maintenance and repopulation defects in the *ddb1*, *atad5a* and *pcna* mutant zebrafish. **A.** Representative images of *cmyb* WISH labeling HSPC population in the CHT at 3 or 4 dpf in *ddb1*^*−/−*^(*cu43/cu43*), *atad5a*^*−/−*^(*cu33/cu33*) and *pcna*^*−/−*^(*cu29/cu29*), and their WT sibling controls. Black brackets indicate *cmyb-*positive HSPCs in CHT. Numbers at the bottom of images indicate the number of representative outcome observed out of the total number of embryos obtained. **B.** Merged images of bright-field and confocal microscopy to show EdU-positive proliferative cells in the CHT between mutants and their WT sibling controls at 3 or 4 dpf and their quantification of the number of EdU-positive cells in the CHT. **C.** Merged images of bright-field and confocal microscopy to display apoptotic cells in the CHT using AO staining between mutants and WT controls at 3 or 4 dpf and their quantification of the apoptotic events in the CHT. Graphs represent mean ± S.E.M. with individual values. p-values were calculated by unpaired two-tailed Student’s t-test. *** p<0.001, **p<0.01. n.s.; not significantly different. Scale bar = 200 μm.

### Sensitivity to genotoxic stress

In contrast to mice, zebrafish loss-of-function mutants of DNA repair genes were mostly viable through embryogenesis. Taking advantage of this unique property, we exposed mutant animals to genotoxic stresses to determine if hypersensitivity to specific types of DNA damage leads to distinct morphology in mutant zebrafish embryos. To induce DNA damage, embryos were treated with ionizing radiation (IR) to induce DNA DSBs or the alkylating agent methyl methanesulfonate (MMS) to stimulate base damage as well as DSBs (Beranek, 1990, Mahaney et al., 2009). Mutant embryos at 6 hpf were exposed to 5 Gy IR or 0.001% MMS and their morphology was screened through 5 dpf (Table S5). Among 30 genes, three mutants (*atad5a*^*−/−*^, *rad51*^*−/−*^ and *shfm1*^*−/−*^) showed hypersensitivity to IR with abnormal morphology in the anterior part of the embryos (Fig. 5A; Fig. S5A). AO staining demonstrated that cell death was induced by IR in the head structures of the *atad5a*^*cu33/cu33*^, *rad51*^*cu54/cu54*^ and *shfm1*^*cu46/cu46*^ mutants (Fig. 5B; Fig. S5B). MMS treatment also induced morphological malformation in the *atad5a*^*−/−*^, *brca2*^*−/−*^, *polk*^*−/−*^, *rad51*^*−/−*^ and *xrcc1*^*−/−*^ mutants (Fig. 5C; Fig. S5C). Apoptosis was robustly induced in the posterior trunk of *atad5a*^*cu33/cu33*^, *brca2*^*cu51/cu51*^, *rad51*^*cu54/cu54*^, and *xrcc1*^*cu59/cu59*^ mutants, while *polk*^*cu13/cu13*^ mutant embryos showed no alteration of apoptotic events in the body trunk where morphological alteration was induced by MMS treatment (Fig. 5D; Fig. S5D). Taken together, our data suggest that seven zebrafish mutants showed hypersensitivity to DNA damaging agents, resulting in induced morphological alterations, which were mainly caused by apoptosis.

**Fig. 5.**
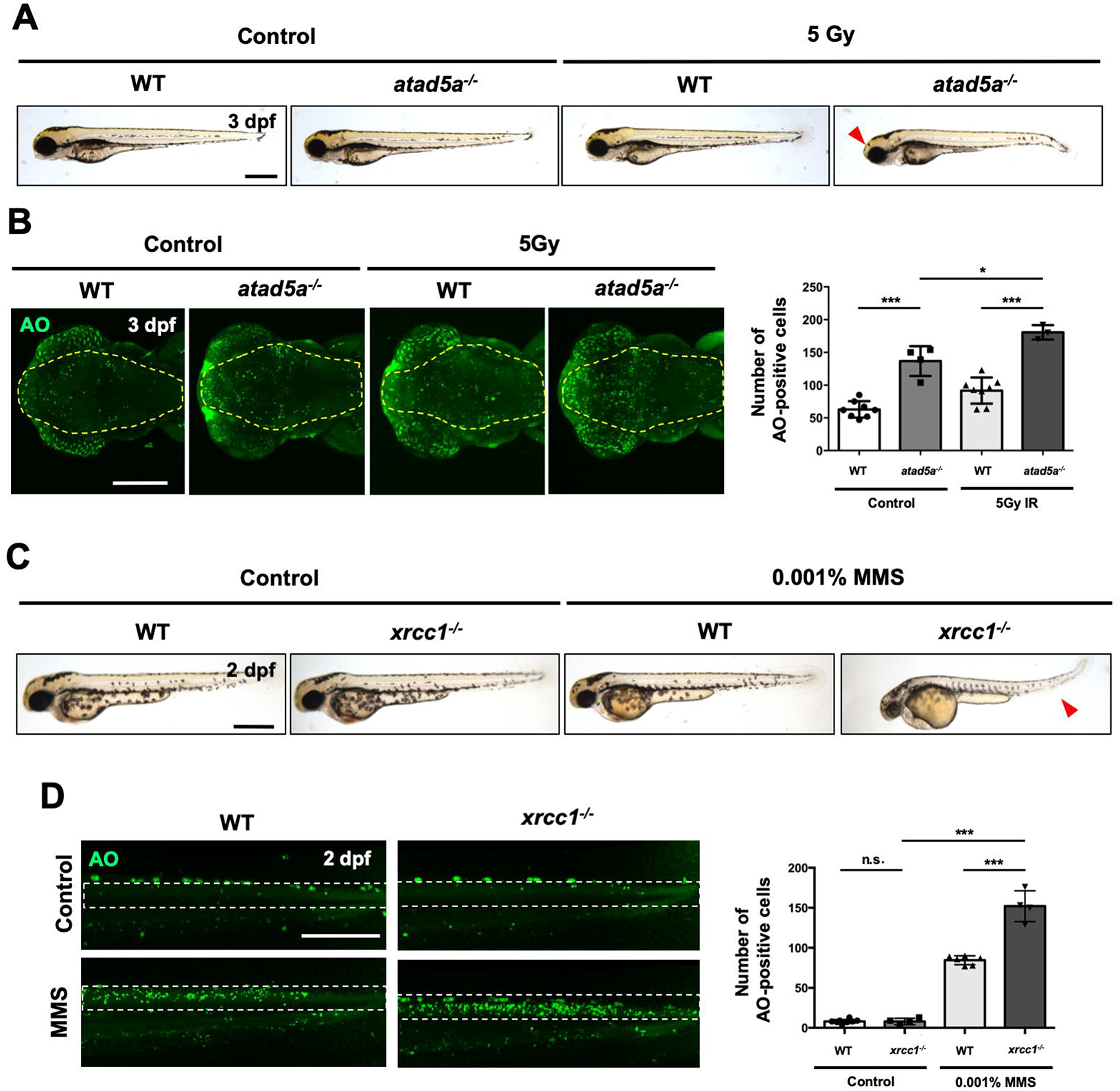
Response of DNA replication/repair mutants to genotoxic stresses. **A.** Images of 3 dpf embryos generated from inbred *atad5a* heterozygous mutant in the presence or absence of 5Gy IR exposure at 6 hpf. Morphological alteration is indicated by red arrowheads in the homozygous mutants exposed to IR. **B.** Confocal microscope images of embryos stained with AO to show apoptotic cells in *atad5a* mutants and WT controls at 3 dpf in response to IR at 6 hpf and its quantification of the number of AO–positive apoptotic cells in the brain of *atad5a*^*−/−*^(*cu33/cu33*) mutants and control with IR-treatment. Yellow dotted area indicates dorsal view of the brain area. **C.** Lateral views of 2 dpf *xrcc1* mutant embryos and WT controls with treatment of 0.001% MMS from 6 hpf. Red arrowheads indicate altered body trunk of the mutants with MMS treatment. **D.** Confocal microscope images showing body trunk of AO-stained 2 dpf *xrcc1*^*−/−*^(*cu59/cu59*) mutant embryos and WT siblings in the presence of 0.001% MMS from 6 hpf and its quantification of AO– positive apoptotic cells in the body trunk of *xrcc1* mutants and WT control with MMS treatment. White dotted box indicates neural tissues in the trunk of embryos. Quantification graphs represent mean ± S.E.M. with individual values. p-values were calculated by unpaired two-tailed Student’s t-test. *** p<0.001, *p<0.05. n.s.; not significantly different. Black scale bar = 500 μm; white scale bar = 200 μm.

### Summary

Homozygous mutants of 18 out of 32 genes resulted in at least one defective in either embryogenesis, growth, sex development, hematopoiesis and hypersensitivity to genotoxic stress (Table 2). In contrast to higher embryonic lethality in KO mice, only 3 out of 32 zebrafish loss-of-function mutants showed lethality before 10 dpf, successfully completing early development, including gastrulation, body patterning and initiation of organogenesis like hematopoiesis. Through further screening of HSPC formation, we found that the same four mutants failed to perform HSPC repopulation during embryogenesis. We discovered additional 13 homozygous mutant animals that displayed abnormal phenotypes at juvenile or adult stages, in an aspect of growth, viability and sex development. Finally, seven mutants showed hypersensitivity to treatment of IR and MMS. Since both damage stimuli can induce DSB, mutants of genes related to HR processes were mainly affected by IR or MMS.

**Table 2.** Summary of mutant phenotyping using 32 CRISPR mutants related to DNA repair processes. Letter “O” indicates defective phenotypes observed in designated mutants with each category of phenotyping.

| DNA repair process | Genes | Phenotyping |  |  |  |  |
| --- | --- | --- | --- | --- | --- | --- |
|  |  | Embryogenesis | Hematopoiesis | Adult survival | Sex reversal | Sensitivity to genotoxic stress |
| BER | *apex1* |  |  | O |  |  |
|  | *mutyh* |  |  |  |  |  |
|  | *nudt1* |  |  |  |  |  |
|  | *unga* |  |  |  |  |  |
|  | *xrcc1* |  |  |  |  | O |
| NER | *ddb1* | O | O |  |  |  |
|  | *ercc1* |  |  |  |  |  |
|  | *neil1* |  |  |  |  |  |
|  | *xpc* |  |  |  |  |  |
| HR | *brca2* |  |  |  | O | O |
|  | *blm* |  |  |  | O |  |
|  | *mre11a* |  |  | O |  |  |
|  | *rad51* |  |  |  | O | O |
|  | *rad54l* |  |  |  | O |  |
|  | *shfm1* |  |  | O |  | O |
|  | *slx1b* |  |  |  |  |  |
|  | *wrn* |  |  | O |  |  |
| TS | *polk* |  |  |  |  | O |
|  | *rad18* |  |  |  |  |  |
| T switch | *hltf* |  |  |  |  |  |
|  | *shprh* |  |  |  |  |  |
| MMR | *msh2* |  |  |  |  |  |
| CLR | *fanci* |  |  |  | O |  |
|  | *usp1* |  |  |  |  |  |
| Replication | *atad5a* | O | O |  |  | O |
|  | *atad5b* |  |  |  |  |  |
|  | *pcna* | O | O |  |  |  |
| Checkpoint | *atrip* |  |  | O |  |  |
|  | *hmgb1a* |  |  |  |  |  |
|  | *ino80* |  |  | O |  |  |
|  | *telo2* |  |  | O |  |  |
| Telomere | *rtel1* |  |  |  | O |  |

### Atad5a is required for normal brain development in zebrafish

We chose the *atad5a* mutants, which showed defects of embryonic morphogenesis and hematopoiesis and displayed hypersensitivity to DNA damage stress, to examine on KO animal in more detail. The ATPase family AAA domain-containing protein 5 (ATAD5) is known for regulating the function of PCNA by unloading PCNA from chromatin during DNA replication or deubiqutinating PCNA following TLS (Kubota et al., 2013, Lee et al., 2010). *Atad5* mutant mice did not survive past embryonic stage before E8.5, implying its crucial role for early development (Bell et al., 2011). Zebrafish possesses two paralogs of ATAD5; Atad5a and Atad5b (Fig. S6A). In our study, we found that loss of *atad5a* leads to developmental aberration including abnormal structure of head and body trunk, and loss of blood stem cell precursors, while loss of *atad5b* displayed no discernable phenotype (Fig. S6B). We further observed that *atad5a* is highly expressed during embryogenesis while the level of *atad5b* was present at much lower levels, suggesting that Atad5a is the main functional paralog in zebrafish (Fig. S6C).

To examine the roles of Atad5a in zebrafish embryogenesis, we analyzed the spatial distribution of *atad5a*. WISH demonstrated that *atad5a* started to express in the whole brain from 1 dpf and specifically localize in the ventricular zone of the midbrain region at 2 dpf, continuing its spatial expression in the brain up to 5 dpf (Fig. 6A; Fig. S6D), suggesting Atad5a is potentially involved in brain development. To further examine the roles of Atad5a on brain development, we first analyzed the functional property of *atad5a*^*cu33/cu33*^ mutants (referred as *atad5*^*−/−*^ for brevity) and observed no functional Atad5a due to nonsense-mediated mRNA decay (NMD) (Tuladhar et al., 2019) (Fig. S7A). Moreover, western blot analysis of whole *atad5a*^*−/−*^ embryo lysates showed that PCNA was accumulated in the chromatin, indicating that PCNA unloading was diminished, similar to what is observed in human cell lines (Kang et al., 2019b) (Fig. 6B). Finally, examination of brain morphology during embryogenesis revealed a clear reduction of brain size in the mutant at 5 dpf (Fig. 6C; Fig. S7B). As no significant difference of whole-body length was found, this suggests a microcephaly-like phenotype of *atad5a*^*−/−*^, indicative of a tissue-specific function (Fig. 6D; Fig. S7C). Taken together, our results demonstrate that Atad5a is important for normal brain development during zebrafish embryogenesis.

**Fig. 6.**
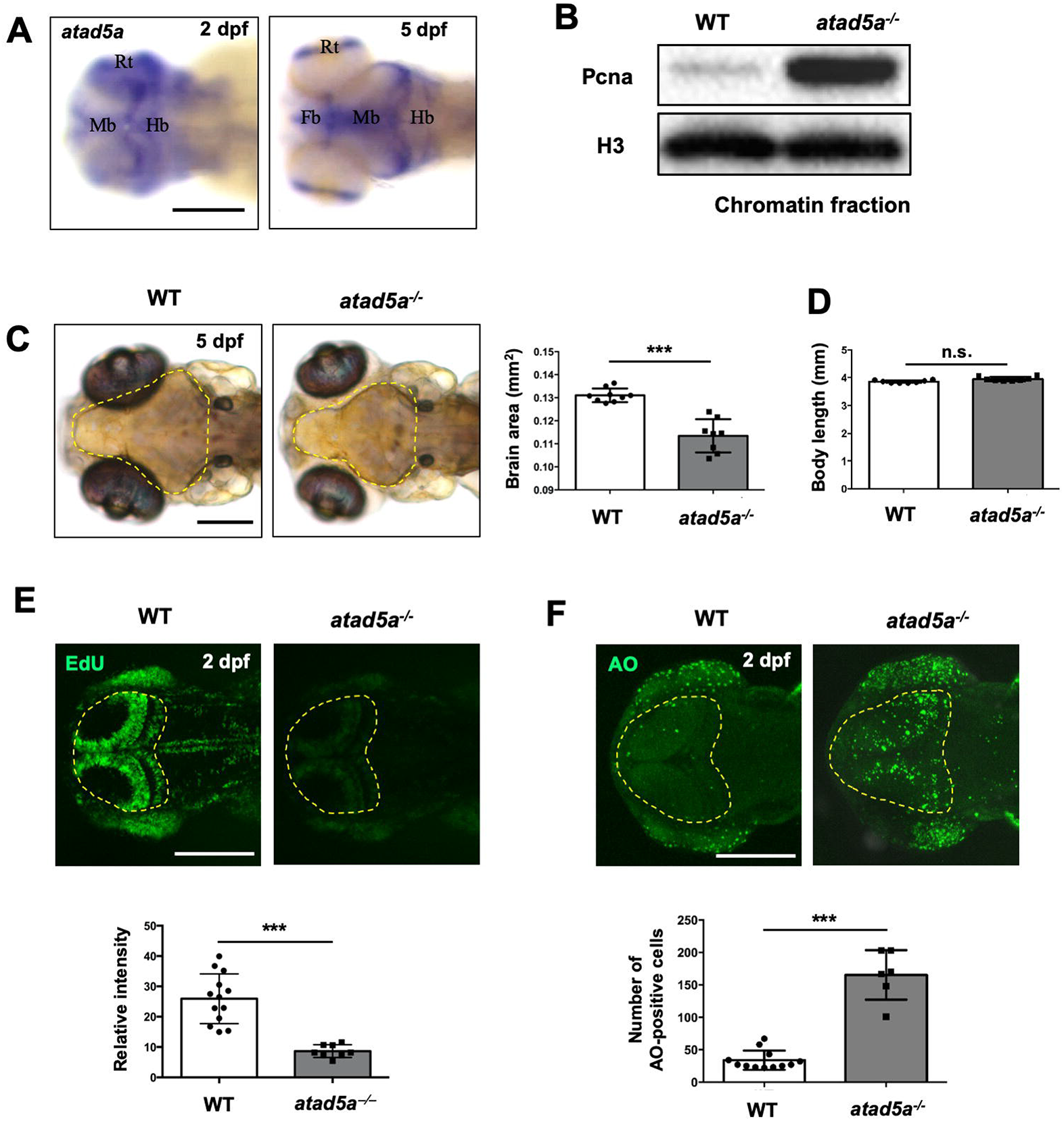
Atad5a is required for brain development during embryogenesis. **A.** Dorsal view of anterior part of zebrafish embryos showing expression of *atad5a* in the brain at 2 and 5 dpf using WISH. *atad5a*-expressing regions are indicated by abbreviated letters; Fb: forebrain, Mb: midbrain, Hb: hindbrain, Rt: retina. **B.** Western blot with Pcna antibody using chromatin fraction of whole embryo lysate from *atad5a*^*−/−*^ and WT controls at 5 dpf. H3 is histone H3 antibody used to show equal loading of chromatin fraction for western blotting. **C.** Dorsal view of anterior part of *atad5a*^*−/−*^ and WT controls at 5 dpf and its quantification of the brain size from dotted area. **D.** Quantification of the entire body length of *atad5a*^*−/−*^ mutants and WT controls at 5 dpf. **E.** Confocal microscope images of EdU-positive cells and quantification of relative intensity of EdU signal in the brain (dotted area) between *atad5a*^*−/−*^ and WT controls at 2 dpf. **F.** Confocal microscope images of AO-positive cells and quantification of AO-positive cell numbers in the brain (dotted area) between *atad5a*^*−/−*^ mutants and WT controls at 2 dpf. Yellow-dotted area indicates the brain including forebrain to cerebellum. All graphs represent mean ± S.E.M. with individual values. p-values were calculated by unpaired two-tailed Student’s t-test. *** p<0.001, **p<0.01, *p<0.05. n.s.; not significantly different. Scale bar = 200 μm.

The ventricular zone, where *atad5a* is expressed, is a highly proliferative region in the developing brain (Hibi and Shimizu, 2012). Moreover, apoptosis was induced by DNA damage in the *atad5a*^*−/−*^ embryos (Fig. 5B). We therefore hypothesized that altered proliferation and/or apoptosis in the brain of *atad5a*^*−/−*^ would lead to malformation of the brain structures of the embryo. We assessed proliferation of brain cells by EdU staining and found EdU-positive cells mainly concentrated in the ventricular zone in WT animals at 2 dpf, while the EdU signal was strongly reduced in the *atad5a*^*−/−*^ mutants from 2 dpf to 5 dpf (Fig. 6E; Fig. S8A). In parallel, AO-positive apoptotic cells were significantly enriched in the mutant at 2 and 5 dpf, compared to WT (Fig. 6F; Fig. S8B). These data suggest that diminished proliferation and induced cell death in the brain of *atad5a*^*−/−*^ embryos leads to abnormal brain formation with microcephaly-like phenotype

### ATM-Chk2 signaling induces p53-dependent apoptosis in the brain of the *atad5a*^*−/−*^ mutants

Along with DNA repair and cell cycle arrest, p53-dependent apoptosis is a well-known consequence of DNA damage induction (Chen, 2016). Interestingly, *p53* transcription was robustly induced in the brain of *atad5a* mutants, compared to WT animals (Fig. 7A). To confirm whether p53-induced apoptosis is responsible for the malformation of the brain, we analyzed the phenotypes of the *atad5a*^*−/−*^; *p53*^*−/−*^ double mutant embryos. Depletion of p53 in the *atad5a* mutant zebrafish reduced apoptosis in the brain and recovered the size of brain without alteration of proliferation, suggesting that the p53 induction in the absence of Atad5a leads to activation of cell death and eventually failure of normal brain development (Fig. 7B, Fig. S9A). To further understand molecular mechanism of p53-depedent apoptotic events, we examined expression of DNA damage markers. Western blot of lysates of whole embryos at 5 dpf showed that γH2AX as well as phospho-Chk2 levels were induced in the *atad5a*^*−/−*^ mutants, suggesting DNA breaks and ATM-Chk2 signaling is activated to induce p53-depndent apoptosis in the absence of Atad5a (Taylor and Stark, 2001) (Fig. 7C). To confirm if ATM activates p53 to induce cell death in the brain without functional Atad5a, we inhibited ATM signaling using the ATM inhibitor KU60019 (ATMi) (Danilova et al., 2014). Perturbation of ATM signaling significantly reduced apoptotic cell numbers in the *atad5a*^*−/−*^ mutants (Fig. 7D). Taken together, our data showed that ATM/Chk2 signaling induces p53-dependent apoptosis in the developing zebrafish brain in the absence of Atad5a.

**Fig. 7.**
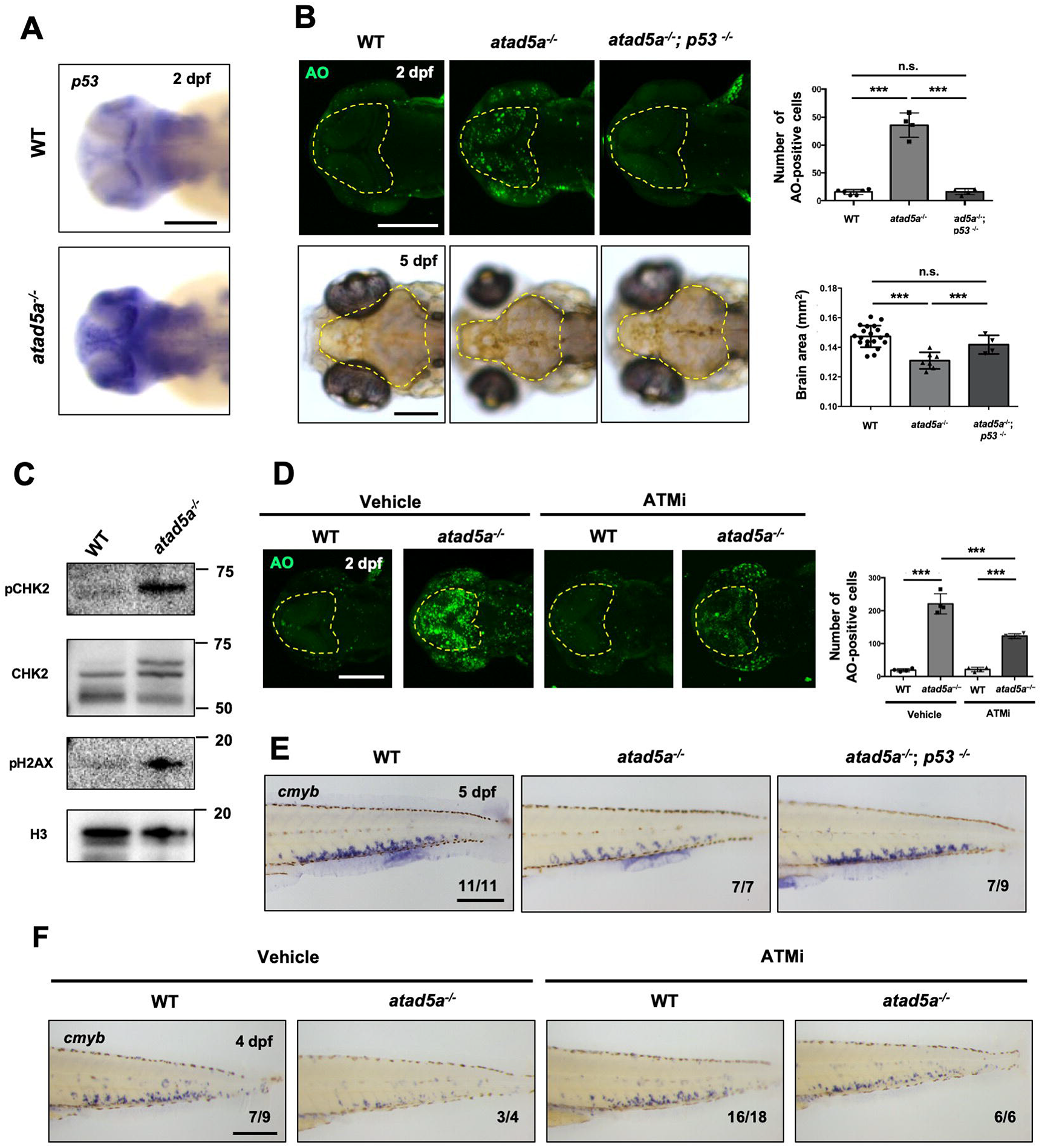
p53-dependet apoptosis in the brain and CHT of *atda5a* mutant is induced by ATM/Chk2 activation. **A.** WISH staining shows that *p53* expression is highly induced in *atad5a* mutant brain compared to WT siblings. **B.** Upper panel shows confocal microscope images of AO-positive cells in the brain (dotted area) of *atad5a*^*−/−*^ single, *atad5a*^*−/−*^; *p53*^*−/−*^ double mutants and WT controls at 2 dpf and their quantification of the number of apoptotic cells in the brain. Bottom panel displays dorsal views of zebrafish head area in the *atad5a*^*−/−*^ single, *atad5a*^*−/−*^; *p53*^*−/−*^ double mutants and WT controls at 5 dpf and quantification of brain size around midbrain and forebrain (dotted area). Yellow-dotted area indicates the brain including forebrain to cerebellum. **C.** Western blot results with antibodies for phospho-Chk2 Thr68 (pChk2), Chk2, and phospho-histone H2AX Ser139 (pH2AX) using whole lysate of WT and *atad5a* mutant embryos at 5 dpf. H3 is histone H3 used to show equal loading of whole lysates for western blotting. **D.** Confocal images of AO staining using WT and *atad5a*^*−/−*^ embryos at 2 dpf with treatment of vehicle control (2% DMSO) and ATMi from 6 hpf and its quantification of the AO-positive cell number in the brain (dotted area). **E.** Images of *cmyb* WISH in the CHT of 5 dpf embryos of *atad5a*^*−/−*^ single, *atad5a*^*−/−*^; *p53*^*−/−*^ double mutants and WT controls. **F.***cmyb* WISH in the CHT of 4 dpf embryos of *atad5a*^*−/−*^ and WT controls with treatment of ATMi from 2 dpf. Numbers at the bottom of images indicate the number of representative outcome observed out of total number of embryos obtained. All graphs represent mean ± S.E.M. with individual values. p-values were calculated by unpaired two-tailed Student’s t-test. *** p<0.001, *p<0.05. n.s.; not significantly different. Scale bar = 200 μm.

### HSPC maturation is blocked by ATM/p53-dependent apoptosis in the *atad5a*^*−/−*^ mutant

Our preliminary investigation of hematopoiesis showed that HSPC precursor cells failed to repopulate in the CHT of the *atad5a* mutants, although HSPC emergence in the dorsal aorta was normal (Fig. 4; Fig. S4B). We found that *atad5a* was expressed in the CHT at 3 dpf, when HSPCs start to repopulate and expand, suggesting its potential requirement for HSPC maintenance and maturation (Fig. S9B). Similar to the brain development, the number of EdU-positive proliferating cells was reduced, while apoptotic events were induced in the CHT of the *atad5a*^*−/−*^ at 3 dpf (Fig. 4). To determine whether p53 activation by ATM/Chk2 signaling induces apoptosis in HSPC precursors in the absence of Atad5a, we performed *p53* WISH assays in the *atad5a*^*−/−*^ and rescue experiment for HSPC marker expression using the *atad5a*^*−/−*^; *p53*^*−/−*^ double mutant and ATMi treatment in the *atad5a*^*−/−*^ animals. *p53* expression was highly induced in the CHT of *atad5a*^*−/−*^ mutants (Fig. S9B). Furthermore, HSPC expansion was rescued in the CHT of *atad5a*^*−/−*^; *p53*^*−/−*^ double mutants as well as ATMi-treated *atad5a*^*−/−*^ embryos, compared to *atad5a*^*−/−*^ single mutants or vehicle-treated *atad5a*^*−/−*^ embryos (Fig. 7E and F). Taken together, our data suggest that *p53* activation by ATM/Chk2 triggered cell death in the HSPC population in the CHT as well as brain of *atad5a* homozygous mutants.

## Discussion

In this study, we generated 32 loss-of-function zebrafish mutants of genes related to DNA repair and replication process, including BER, NER, HR, TLS and MMR, via multiplexed CRISPR mutagenesis followed by florescence PCR-based fragment analysis (Varshney et al., 2015). To fully understand functional annotations of genes *in vivo*, we screened the phenotypes of the mutants in embryogenesis, viability, growth, sex determination, hematopoiesis and sensitivity to DNA damaging agents. All the 32 homozygous mutants grew normally until 48 hpf with only four mutants showing morphological malformation and hematopoietic defects before 6 dpf, in line with observations that genomic homeostasis is linked to hematopoiesis (Botthof et al., 2017, Marsh et al., 2018, Farres et al., 2015). Mutants of *pcna* and *atad5a*, two critical genes for DNA replication (Kang et al., 2019a), showed reduced proliferation and induced apoptosis in the blood precursors at the CHT, while *ddb1* homozygous mutants showed loss of HSPCs caused by reduced proliferation. PCNA and ATAD5 are critical for not only general DNA replication but also for the maintenance of replication fork stability (Park et al., 2019). Perturbation of genomic stability generally induces diverse DNA damage responses, such as cell-cycle arrest and cell death (Jiang et al., 2017, Gao et al., 2018, Wang et al., 2016). Activation of ATM pathway and p53-apoptotic process in the *atad5a* zebrafish mutants suggest that maintenance of genomic stability is critical for maintenance and repopulation of HSPCs.

Further monitoring viability and development of the mutants demonstrated that seven more KO zebrafish showed defects of survival during larval and juvenile stages; *shfm1*^*−/−*^, *telo2*^*−/−*^, *apex1*^*−/−*^, *atrip*^*−/−*^, *mre11a*^*−/−*^, *wrn*^*−/−*^ and *ino80*^*−/−*^. Unlike KO studies using murine system, all the mutant animals showed distinct defects of morphological appearance or growth at different developmental stages after embryogenesis. While KO mouse studies for the function of *Shfm1* is not yet reported, systematic analysis of KO mice in the other 6 genes have shown their critical roles in early embryogenesis resulting in embryonic lethality (Takai et al., 2007, Xanthoudakis et al., 1996, Dickinson et al., 2016, Luo et al., 1999, Lombard et al., 2000, Oshima et al., 1996). Conditional KO (CKO) mouse approaches have shown the roles of Apex1 on neurological function with injury and functions of Ino80 on genomic stability in mammals (Min et al., 2013, Stetler et al., 2016). In addition, our six zebrafish mutants showed female-to-male sex reversal defects; *blm*^*−/−*^, *brca2*^*−/−*^, *fanci*^*−/−*^, *rad51*^*−/−*^, *rad54l*^*−/−*^ and *rtel1*^*−/−*^. While *rad54l*^*−/−*^ and *rtel1*^*−/−*^ mutants demonstrated a possible sex reversal phenotype with normal fertility like *fanci*^*−/−*^ homozygotes (Ramanagoudr-Bhojappa et al., 2018), *blm*^*−/−*^ male animals were infertile without normal functional germ cells in the testis like *brca2*^*−/−*^ and *rad51*^*−/−*^ where primordial germ cells failed to survive and proliferate in the testes (Botthof et al., 2017, Shive et al., 2010). Although *Blm* Homozygous KO mouse embryos were developmentally delayed and died by embryonic day 13.5 (Chester et al., 1998), *blm*^*cu53/cu53*^ zebrafish survived and we were able to observe aberrant spermatogenesis, potentially recapitulating a phenotype of human male infertility in patients with Bloom syndrome (de Renty and Ellis, 2017). Taken together, our zebrafish KO mutants showed diverse developmental problems at various stages, implying distinct roles of each gene *in vivo*. Further phenotypic annotation of each zebrafish mutant generated in our study will provide valuable knowledge of biological roles of DNA repair genes excepting early embryogenesis without laborious CKO approaches.

Our CRISPR mutants showed phenotypes consistent with previously reported zebrafish loss-of-function studies, including the sex reversal defects of *brca2*^*−/−*^, *rad51*^*−/−*^ and *fanci*^*−/−*^ mutants (Shive et al., 2010, Botthof et al., 2017, Ramanagoudr-Bhojappa et al., 2018) and embryonic malformation with aberrant head development in *ddb1*^*−/−*^ zebrafish embryos (Hu et al., 2015). Interestingly, several zebrafish knock-down studies using morpholinos (MOs) yielded different results from our mutant studies. For example, knock-down of BER components, *apex1* or *unga* using MOs led to aberrant heart and brain development during embryogenesis (Wang et al., 2006, Pei et al., 2019, Wu et al., 2014) and *hmgb1a* morphants showed abnormal morphology of the brain during early development (Zhao et al., 2011), whereas mutants of these three genes develop normally in embryogenesis. This discrepancy between studies of MOs and gene-edited mutants could be due to several reasons. First, early knock-down using translation blocking MOs might block the gene function earlier than KOs by blocking expression of target genes in maternal mRNA, removing maternal protection and resulting earlier phenotypes. To clarify this hypothesis using mutant animals, phenotyping maternal zygotic mutant embryos will be practical. The second possible cause is a potential genetic compensation in the zebrafish CRISPR mutants, which is rarely observed in the stable KO models by inducing genes related to the mutated one (Rossi et al., 2015). The presence of genetic compensation phenomenon can be identified by transcriptome analysis between the mutants and their sibling controls.

We further examined sensitivity of the mutants to two genotoxic agents. IR generally induces DNA DSBs, while MMS is known as an alkylation agent that triggers DNA repair processes including DSB and excision repair as it forms alkylated bases and DSBs (Beranek, 1990, Mahaney et al., 2009, Lundin et al., 2005). As expected, mutants of HR-related genes like *rad51* and *shfm1* showed hypersensitivity to DSBs induced by IR, while *xrcc1*^*−/−*^ mutants with an BER defect were sensitive to MMS treatment. Additionally, MMS-induced hypersensitivity in the mutants of *polk* essential for TLS process is consistent with the study of *C. elegans polk* null mutants where mutation rate in the genome is robustly enhanced by MMS treatment (Volkova et al., 2020). Likewise, MMS-induced malformation of our zebrafish *polk*^*−/−*^ mutants without induced apoptosis is possibly caused by highly induced mutagenesis in the somatic genome of the mutant. This is consistent with the role of Polk in bypassing minor groove DNA adducts such as 3-methyladenine (Plosky et al., 2008). In addition to known function of each DNA repair gene, we uncovered potentially unknown functions of genes like *atad5a*. Beside 18 mutants with defects of development or genotoxic sensitivity (Table 2), the other 14 zebrafish mutants showed no altered phenotypes in our assays. To closely dissect these mutant animals without visible phenotypes in this study, it is important to prioritize more specific phenotyping strategies based on their general cellular functions. Therefore, further investigation of sensitivity to genotoxic stress using various DNA damage agents may lead to observation of new morphological alterations. In addition, further screening of non-lethal tissue-specific phenotypes along with analysis of genome expression will provide more information of functional roles of these DNA repair genes in a vertebrate.

To successfully complete highly proliferative neurogenesis, maintenance of genomic stability and DNA damage signaling are both critical (McKinnon, 2009, McKinnon, 2013). Loss of genomic stability by malfunction of DNA repair protein often leads to neurologic disease (Bender et al., 2006, Reynolds et al., 2009). Consistent with previous cellular studies, we found that *atad5a* mutant zebrafish showed sensitivity in response to both IR and MMS (Lee et al., 2013, Park et al., 2019). In addition, chromatin-bound PCNA is accumulated in the mutant background, showing that the cellular function of ATAD5, the removal of PCNA upon completion of replication, is conserved in zebrafish. *atad5a*^*−/−*^ embryos showed reduced size of the brain where proliferation was diminished while cell death was induced by ATM/p53 activation. Based on our characterization of *atad5a*^*−/−*^ phenotype, we speculate loss of Atad5a leads to the accumulation of PCNA on chromatin, causing replication stress and induction of cell death through ATM/Chk2 and p53. ATM is one of main DNA damage-activated kinases in the nervous system, responding to mainly DSB (Shiloh and Ziv, 2013). Our data describing the function of *atad5a* using zebrafish mutants demonstrated that Atad5 plays a pivotal role to maintain genomic integrity and regulate DNA damage signaling for normal neurogenesis and brain development.

Through high-throughput characterization of genetically functional materials, zebrafish KO lines can be utilized for functional studies of specific proteins with emphasis on the organ level or systemic phenotype on a large scale. Survival/lethality and phenotypic studies of DNA damage response genes will not only help unravel the importance of those genes at specific tissues, but also show the functional spectrum of influence on the disease phenotypes. These model systems can therefore also be vital platform technology for the study of genetic therapy and drug testing for human diseases.

## ACKNOWLEDGEMENT

We thank Hana Kim and Yun Ji Choi for technical assistance, Orlando Sharer for scientific guidance and members of Molecular Genetics Section in the Center for Genomic Integrity for providing helpful comments.

## Author Contribution

Conceptualization and Methodology, Y.L. and K.M.; Validation, U.S., K.N., C.O. and Y.L.; Formal Analysis and Visualization, U.S., K.N., C.O., and Y.L; Investigation, U.S., K.N., C.O., B.C., H.S., G.V., Y.K., H.S., S.J., G.R., S.K., S.Y., Y.C., and S.P.; Writing-Original Draft, U.S., K.N., C.O. and Y.L.; Supervision, Y.L. and K.M.; Project Administration, Y.L. and K.M.

## DATA AVAILABILITY

## Materials and methods

### Zebrafish husbandry

All zebrafish embryos and adult animals were raised and maintained in a circulating aquarium system (Genomic Design or Aquaneering) at 28.5◻°C in accordance with UNIST or NIH Institutional Animal Care and Use Committees (IACUC: UNISTIACUC-20-09; NIH G-05-5). For CRISPR mutant generation, wild-type TAB5 strain was used. For loss of function studies for p53, *tp53*^*zdf1/*zdf1^ line was used (Berghmans et al., 2005).

### Generation of zebrafish mutants through CRISPR/Cas9 mutagenesis

Single guide RNAs (sgRNAs) targeting 57 genes related to DNA repair processes were designed and generation of sgRNA template and preparation of sgRNA and *Cas9* mRNA was performed as previously described (Varshney et al., 2015, Varshney et al., 2016) (Supplementary Table S1). Up to four gRNAs (50pg per each gRNA) targeting different genes mixed with *Cas9* mRNA (300pg) were injected into wild-type TAB5 embryos at 1-cell stage grown to adulthood. The multiplexing scheme and fluorescence PCR primer sequences for each gene are described in Supplementary Table S1. To confirm germline-transmitted in/del mutations, F1 progeny generated from injected founder fish were screened through fluorescence PCR as described previously (Varshney et al., 2016). Sanger sequencing was performed to determine the sequence of mutant alleles.

### Mutant genotyping

Genomic DNA extractions were performed with Extract-N-Amp or REDExtract-N-Amp™ Tissue PCR Kits (Sigma-Aldrich). Whole embryos or adult zebrafish fin-clips were lysed by incubation in a mixture of 50 μL extraction solution (Extraction Solution E7526) and 14 μL tissue preparation solution (Tissue Preparation Solution T3073) at room temperature for 10 min, 95°C for 5 min, 25°C for 5 min and then neutralized with 50 μL of neutralization solution (Neutralization Solution B N3910). Aliquots from these lysates were used for a PCR (with a Taq DNA polymerase with standard Taq buffer NEB or EconoTaq DNA polymerase Lucigen) with respective gene primers and an 18mer M13F-FAM fluorescent tagged primer (5’-TGTAAAACGACGGCCAGT-3’) in a PCR condition: 94°C (12min) denaturation, 35 cycles of amplification (94°C for 30 sec, 57°C for 30 sec, 72°C for 30 sec), and a final extension at 72°C for 10 min, and indefinite hold at 4°C. Genetic fragment analyses to identify mutant amplicons generated by insertion or deletion were performed on a 3130xI Genetic Analyzer (Applied Biosystems Hitachi) and the data was analyzed in a GeneMapper v.4.0 as per protocol described (Varshney et al., 2015).

### Mutant phenotyping

Heterozygous adult zebrafish were in-crossed and the clutches were examined under a Leica Stereo-Microscope (M80) daily until 7 days post fertilization (dpf) to monitor potential embryonic defects. The clutches were imaged on a LEICA M165C digital stereo microscope during early developmental stages. For the mutants with normal embryogenesis, the clutches were grown to around four months and the alleles for the adult survival screens were assayed with genetic fragment analysis. To measure the growth of the mutants and their sibling controls, fork lengths (from the snout to the end of the middle of the caudal fin) were measured for each progeny and then sampled for genotype. A student t-test was performed for the plotted body sizes for each mutant assayed. The parameters were set for an unpaired t-test with two-tailed P values set for a confidence interval at 95% (GraphPad Prism 5).

### Fertility test

The homozygote males showing potential sex reversal phenotype (*blm*^*−/−*^, *brca2*^*−/−*^, *fanci*^*−/−*^, *rad51*^*−/−*^, *rad54l*^*−/−*^, *rtel1*^*−/−*^) were selected and individually mated to WT females. Minimum of 8 outcrosses were made from each mutant in separate mating tanks. All eggs from each hatched clutch were collected and incubated at 28°C and fertilization screening was performed at 24 hpf. Genotyping was performed using ≤16 fertilized eggs per clutch to validate the parents’ genotype (−/−male x +/+ female).

### Western blotting and immunoprecipitation

Protein samples were prepared from zebrafish embryos (70 embryos of wild-type and *atad5a* mutant at 5 dpf) or cells as either soluble/chromatin fraction or whole lysate. For soluble/chromatin faction, samples were lysed first using Buffer A (100 mM NaCl, 300mM Sucrose, 3 mM MgCl_2_, 10 mM PIPES, 1 mM EGTA and 0.2% Triton X-100) containing the phosphatase inhibitor PhosSTOP (Roche) and complete protease inhibitor cocktail (Roche)) for 8◻min on ice. Crude lysates were centrifuged at 5000◻×◻rpm, 4◻°C for 5◻min to separate the chromatin-containing pellet from the soluble fraction. Remaining pellet was lysed using RIPA buffer (50◻mM Tris–HCl (pH 8.0), 150◻mM NaCl, 5◻mM EDTA, 1% Triton X-100, 0.1%◻SDS, 0.5% Na deoxycholate, 1◻mM PMSF, 5◻mM MgCl_2_) containing the phosphatase inhibitor PhosSTOP, complete protease inhibitor cocktail and benzonase. The chromatin fraction was clarified by centrifugation at 13,000◻×◻rpm, 4 °C for 5◻min. For immunoprecipitation, whole-cell lysates were prepared by lysing cells with buffer X (250◻mM NaCl, 5◻mM MgCl_2_, 100◻mM Tris–HCl (pH 8.5), 1◻mM EDTA, and 1% Nonidet P-40) containing the phosphatase inhibitor PhosSTOP, complete protease inhibitor cocktail, and 500 units of Benzonase. Lysates were cleared by centrifugation at 13,000◻×◻rpm, 4◻°C for 5◻min. For HA or Myc immunoprecipitation, HA-tagged or Myc-tagged proteins were isolated with anti-HA agarose or anti-c-Myc Agarose Affinity Gel (Sigma), respectively. Anti-HA and anti-Myc agarose beads were resuspended in 1× SDS loading buffer. Following antibodies were used in the experiments with indicated dilution; anti-Pcna (GENETEX GTX124496, 1:1000), anti-PCNA (PC10) (Santa Cruz Biotechnology, sc-56, 1:2000), anti-histone H3 (upstate 07-690, 1:15000), anti-HA (HA-7) (Sigma, H-9658, 1:10000), anti-CHK2 (Abcam, ab8108; 1:1000), anti-pCHK2 (T68) (Cell Signaling Technology; #2197; 1:1000), anti-pH2AX (S139) (Cell Signaling Technology; #9718; 1:1000), anti-ATAD5 (Lee et al., 2013) (1:2000)

### Whole mount in situ hybridization (WISH)

Embryos were fixed using 4% paraformaldehyde for overnight at 4°C. Fixed embryos were washed using PBS and dehydrated in methanol at −20°C for overnight. After dehydration, embryos were permeabilized using acetone at −20°C. Samples were hybridized with a digoxigenin (DIG)-labelled antisense RNA probe in hybridization buffer (50% formamide, 5xSSC, 500◻μg/ml Torula yeast tRNA, 50◻μg/ml heparin, 0.1% Tween-20 and 9◻mM citric acid (pH 6.5)) for 3days at 65°C. After hybridization, samples were washed with 2xSSC solution and 0.2xSSC solution. Washed samples were incubated with alkaline phosphate-conjugated DIG antibodies (1:4000) (Roche, Mannheim, Germany) for overnight at 4°C. Samples were developed using alkaline phosphate reaction buffer (100◻mM Tris, pH9.5, 50◻mM MgCl_2_, 100◻mM NaCl and 0.1% Tween-20) with the NBT/BCIP substrate (Promega) to visualize the WISH signal.

### 5-Ethynyl-2’deoxyuridine(EdU) assay

Zebrafish embryos were saturated in 10 mM EdU in E3 fish water with 15% DMSO for 10 min. Embryos were fixed with 4% paraformaldehyde in PBS. Fixed embryos were dehydrated in methanol at −20°C. Embryos were then rehydrated in PBS with 0.1% Tween-20 (PBT) and penetrated with 1% Triton X-100. EdU signal was detected using Click-iT EdU Alexa Fluor 488 Imaging Kit (Invitrogen) and samples were imaged using LSM880 confocal microscopy (Carl Zeiss).

### Acridine orange (AO) stain

Embryos were incubated in 20 μg/mL AO solution dissolved in E3 water for 10 min. After washing twice with E3 fish water, samples were immediately imaged with LSM880 confocal microscope.

### Ionizing radiation and chemical treatment

To induce DNA DSBs, 5Gy IR was given to zebrafish embryos collected at 6 hours post fertilization (hpf) from heterozygous in-crosses using Gammacell® 3000 Elan (Best Theratronics). Irradiation time for 5 Gy was determined depending on the decay factor of Caesium-137 source. For methyl methanesulfonate (MMS) treatment, embryos were continuously incubated from 6 hpf in E3 water with 0.001% MMS. Embryos were examined for possible phenotypes under a Leica Stereo-Microscope (M80). For ATM inhibitor (ATMi) treatment, embryo clutches generated from *atad5a* heterozygous in-crosses were incubated with 200 μM ATMi (KU60019; Sigma-Aldrich) or 2% DMSO dissolved in E3 fish water from 6 hpf until 48 hpf for the examination of brain development and from 2 to 4 dpf for hematopoiesis.

### Histological assay

Testes dissected from adult zebrafish were immediately fixed in 4% paraformaldehyde at 4°C for 24 hours and then incubated in a 10% NBF (neutral buffered formalin) at 4°C for 24 hours. The tissues were then paraffin-embedded and sectioned into 4 μm thickness by microtome (Leica RM2255) on a coating slide. After deparaffinization with Xylene (Samchun, X0020), hydrated slides were stained with hematoxylin (Mayer’s: YD-diagnostics, S2-5) and rinsed with water. The slides were then processed routinely in HCl and ammonia water. Excess reagent was rinsed off and counterstained in Eosin (Duksan, c73521). The slides were dehydrated from 70% EtOH to 100% EtOH, rinsed in xylene, and mounted with mounting solution (Muto, 2009-2).

### Quantitative real-time RT-PCR

Total RNA was extracted from zebrafish embryos with 1 ml of TRIzol and 0.2 ml of chloroform and precipitated with 0.5 ml of isopropanol. One microgram of RNA was then reverse transcribed with SuperScript Reverse Transcription kit (Invitrogen). Quantitative real-time PCR was performed using Power SYBR Green Master Mix (Applied Biosystems). Temperature profile for the reaction was 50°C for 2 min, 95 °C for 3 min, and then 95°C for 15 s and 60°C for 30 s for 55 cycles. The expression levels were relatively quantified using comparative threshold method and normalized to βactin as endogenous control for *atad5a* and *atad5b*. Following primers were used for this experiments; *atad5a*_forward-5’-GCGCCACAGAAGAAAGAGAA-3’, *atad5a*_reverse-5’-GTTGGAGGCATTCGACTGAT-3’, *atad5b*_forward-5’-GAGCAGACACCGAAGAGAGG-3’, *atad5b*_reverse-5-AGCTCTTGTGCACAGGCATA-3’, β*-actin*_forward-5’-CGAGCTGTCTTCCCATCCA-3’, β*-actin*_reverse-5’-TCACCAACGTAGCTGTCTTTCTG-3’.

### Cell culture

Human embryonic kidney (HEK) 293T cells were cultured in Dulbecco’s modified Eagle’s medium (Hyclone) supplemented with 10% fetal bovine serum (Hyclone) and 1% penicillin-streptomycin (Gibco) at 37◻°C under 5%◻CO_2_. Plasmids expressing wild-type PCNA or its mutants were cloned into pcDNA3 mammalian expression vector. All constructs were confirmed by sequencing. Plasmid DNA was transfected to HEK293T cells using X-tremeGENE HP DNA transfection Reagent (Roche). Transfected cells were harvested at 48◻hours after transfection for further analysis.

