## Supplemental Data for "High-throughput generation and phenotypic characterization of zebrafish CRISPR mutants of DNA repair genes"

**Supplemental Figures and Tables**

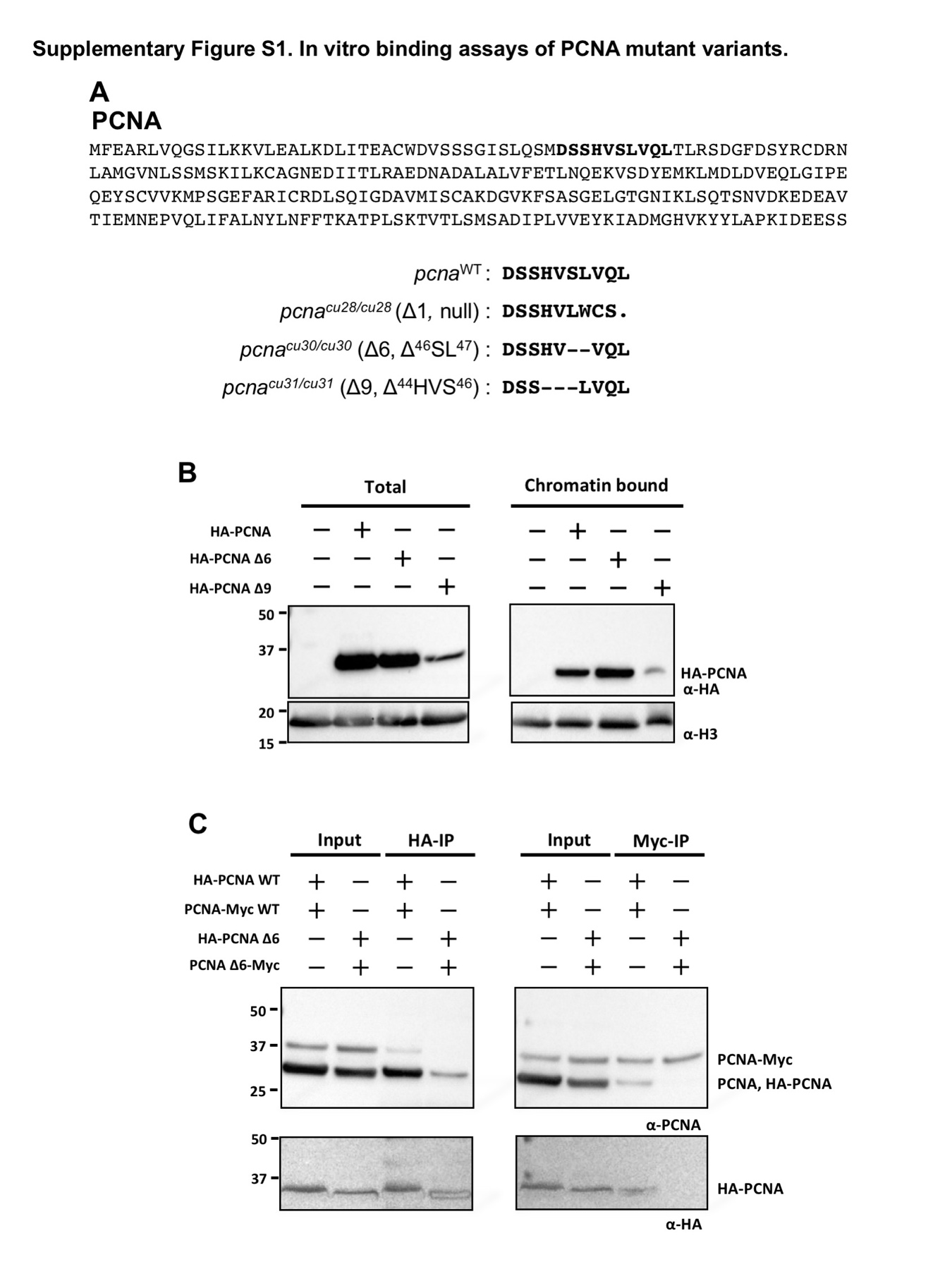

**Supplementary Fig. S1. *in vitro* binding assays of PCNA mutant variants. A.** Whole amino acid sequence of zebrafish PCNA and the sequences of sgRNA target site between wild-type and *pcna* mutants including *pcna^cu28/cu28^* null mutant and inframe mutants like *pcna^cu30/cu30^* and *pcna^cu31/cu31^*. Bold letters indicate sgRNA-targeting amino acid sequences. A period indicates a premature stop in *pcna^cu28/cu28^* null mutants, while hyphens indicate deleted sequences of *pcna^cu30/cu30^* and *pcna^cu31/cu31^* mutants. **B.** Immunoblot analysis with total or chromatin bound PCNA. HA-tagged wild-type or mutant PCNA were transiently expressed in human HEK293T cells. Expression of PCNA Δ9 mutant is low, while PCNA Δ6 and wild-type PCNA were expressed normally in the whole cells or chromatin-bound forms of human HEK293T cells. **C.** PCNA Δ6 was defective in making hetero-complex with wild-type PCNA. Myc tagged wild-type or Δ6 mutant PCNA was co-expressed with HA tagged wild type or Δ6 mutant HA-PCNA in HEK293T cells. PCNA was immunoprecipitated from the lysate using anti-HA(left) or anti-Myc(right) antibody conjugated agarose bead. Co-precipitated, differently tagged PCNA were analyzed by immunoblot. HA tagged PCNA was overlapped to endogenous PCNA on immunoblot, so amount of HA tagged PCNA was analyzed by anti-HA tag antibody.

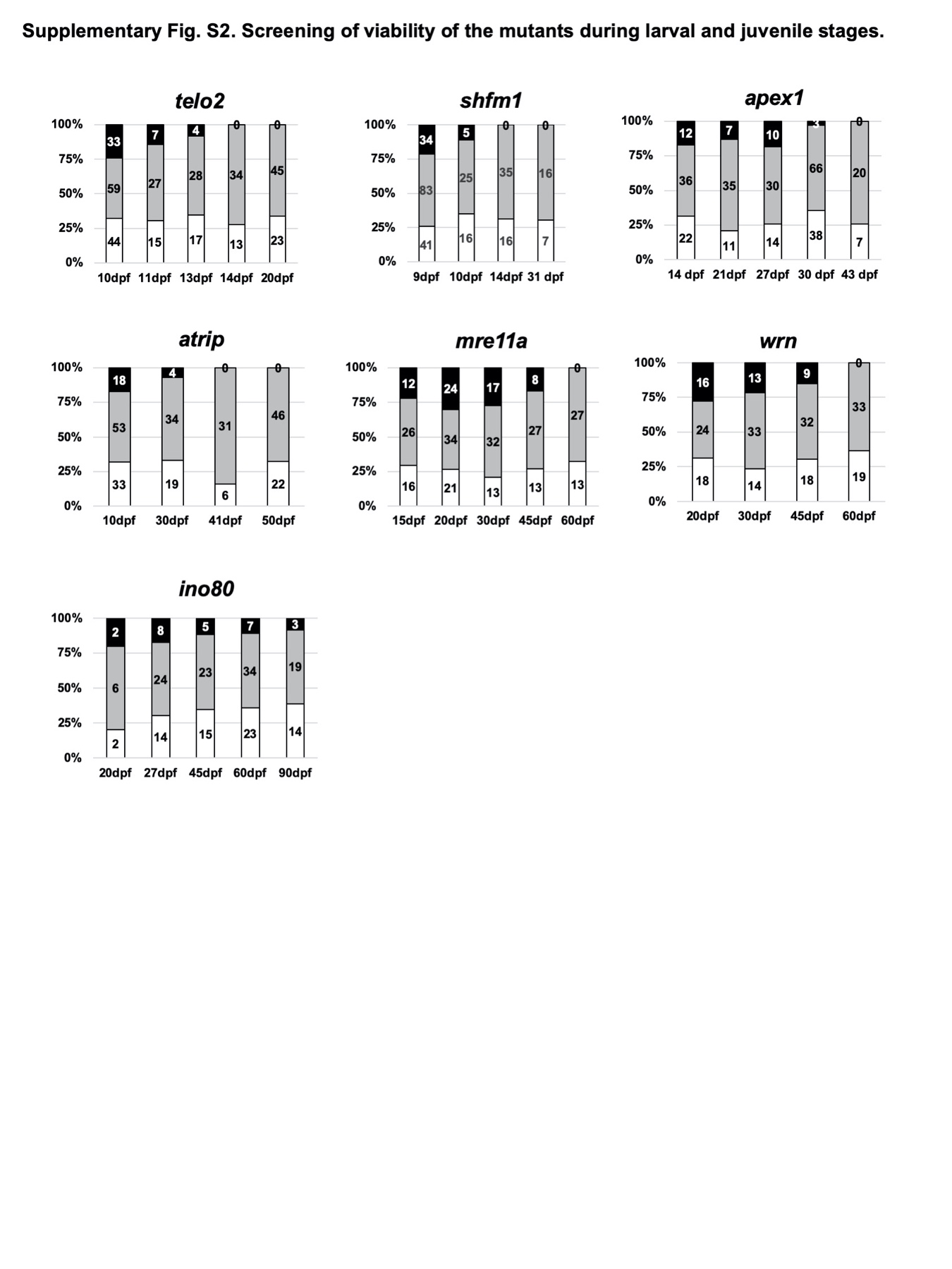

**Supplementary Fig. S2. Screening of viability of the mutants during larval and juvenile stages.** The number of genotyped zebrafish progeny from inbred heterozygous mutants of *telo2*, *shfm1*, *apex1*, *atrip*, *mre11a*, *wrn* and *ino80* at designated developmental stages with three representative genotypes (+/+, +/-, -/-). Numbers in the bar of each graph represent the number of fish with each genotype.

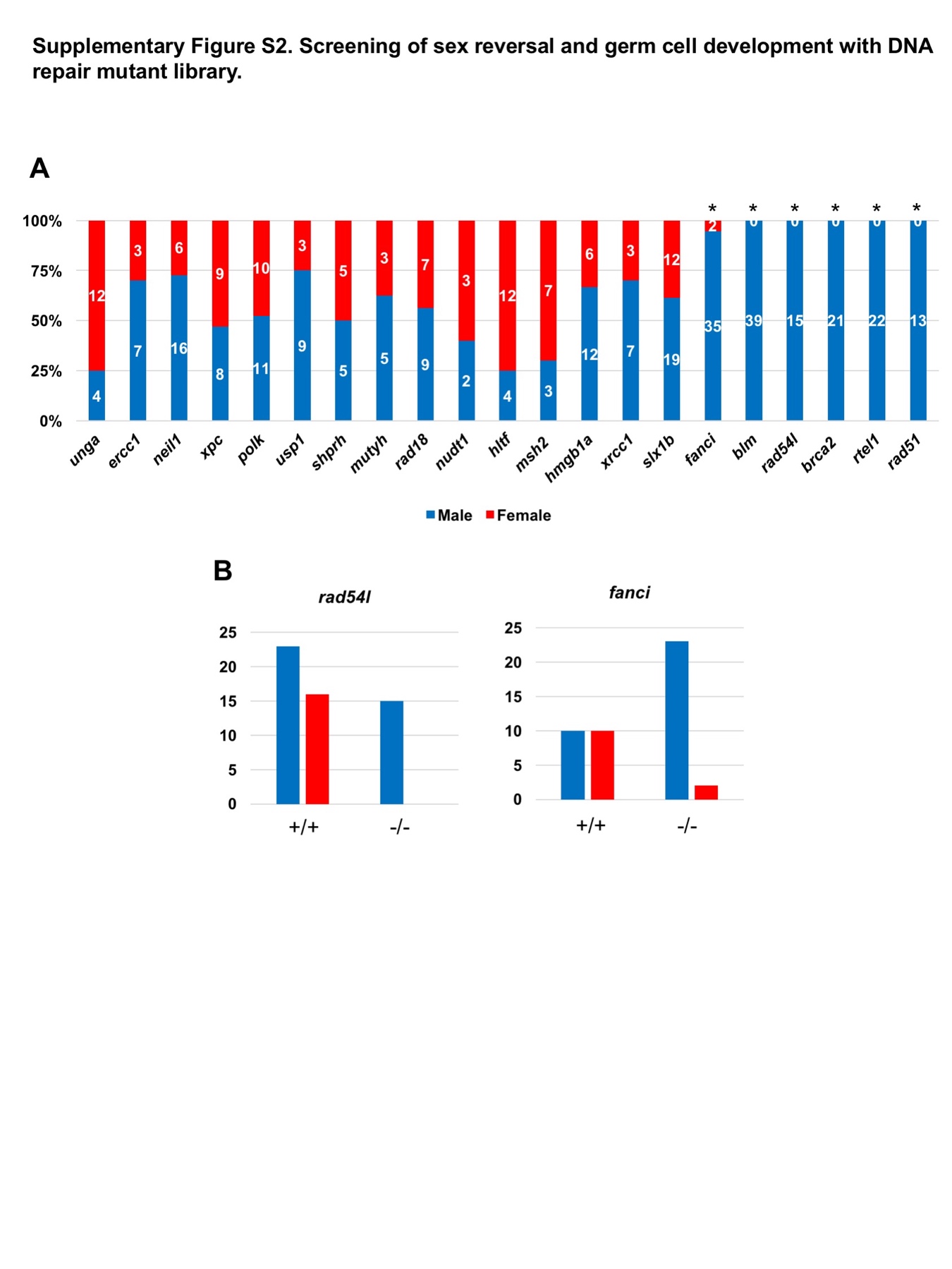
**Supplementary Fig. S3. Screening of sex reversal and germ cell development with DNA repair mutant library. A.** Sex of 21 homozygous mutants of DNA repair genes was determined at 4-9 month old. Asterisks indicate homozygotes which failed to generate female fish possibly due to sex reversal. **B.** The number of male and female animals genotyped (+/+ and -/-) from inbred heterozygous mutants of *rad54l and fanci.*

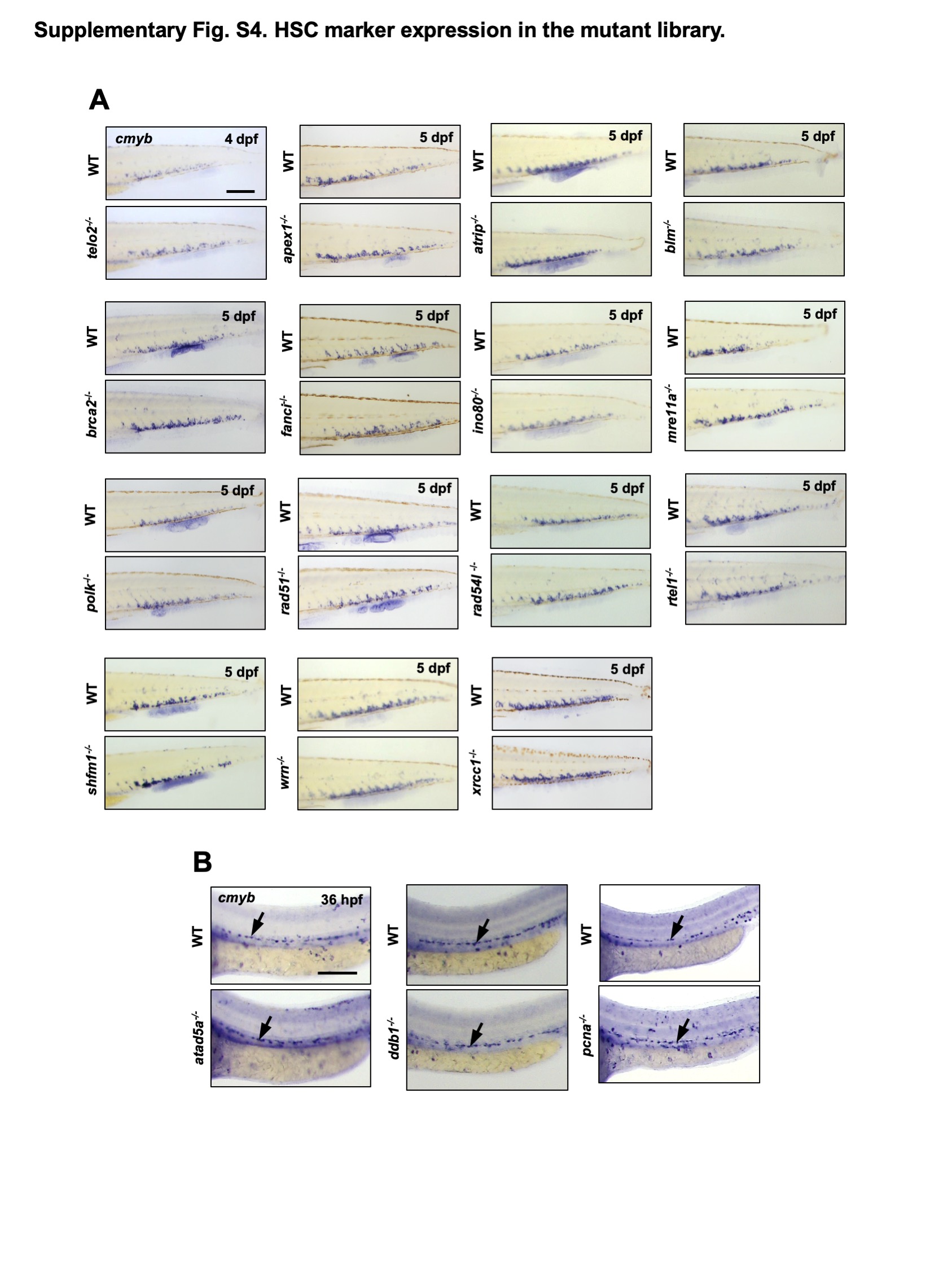
**Supplementary Fig. S4. HSC marker expression in the mutant library. A.** Screening of HSC marker expression in the CHT using *cmyb* at 4 or 5 dpf using 15 mutants and WT sibling controls; *telo2^-/-^(cu41/cu41), apex1^-/-^(cu2/cu2)*, *atrip^-/-^(cu37/cu37), blm^-/-^(cu53/cu53),* *brca2^-/-^*(*cu51/cu51*), *fanci^-/-^*(*cu24/cu24*), *ino80^-/-^(cu36/cu36), mre11a^-/-^(cu56/cu56),* *polk^-/-^*(*cu13/cu13*), *rad51^-/-^*(*cu54/cu54*), *rad54l^-/-^*(*cu62/cu62*), *rtel1^-/-^(cu42/cu42), shfm1^-/-^(cu46/cu46), wrn^-/-^(cu64/cu64). xrcc1^-/-^*(*cu59/cu59*)*. cmyb* expression was not altered in these 15 mutants, compared to WT sibling controls. **B.** Images of *cmyb* WISH labeling HSPCs emerging from dorsal aorta at 36 hpf (black arrows), using embryos of *atad5a^-/-^*(*cu33/cu33*), *ddb1^-/-^*(*cu43/cu43*) and *pcna^-/-^*(*cu29/cu29*), and their WT sibling controls. Scale bar = 200 µm

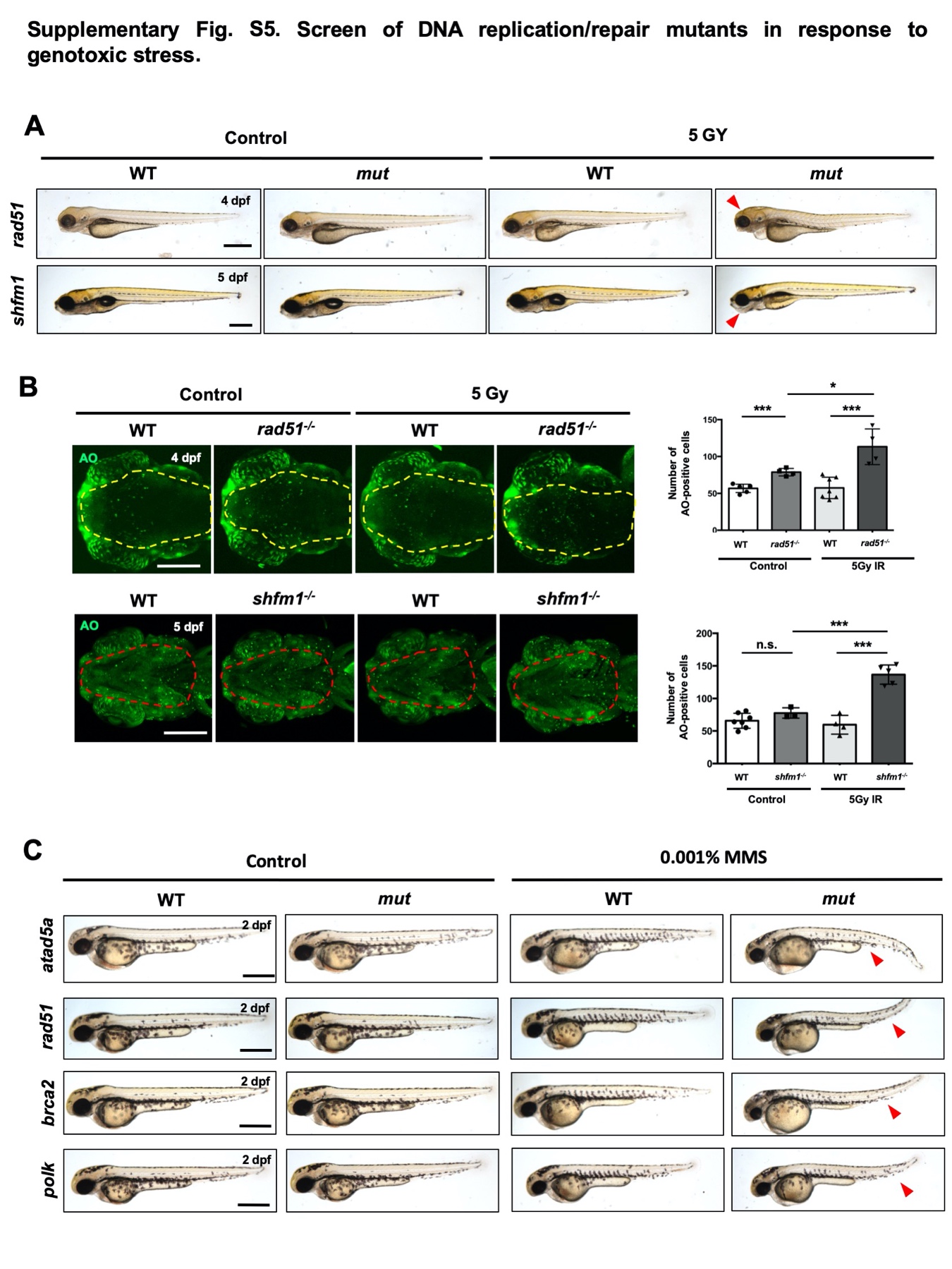

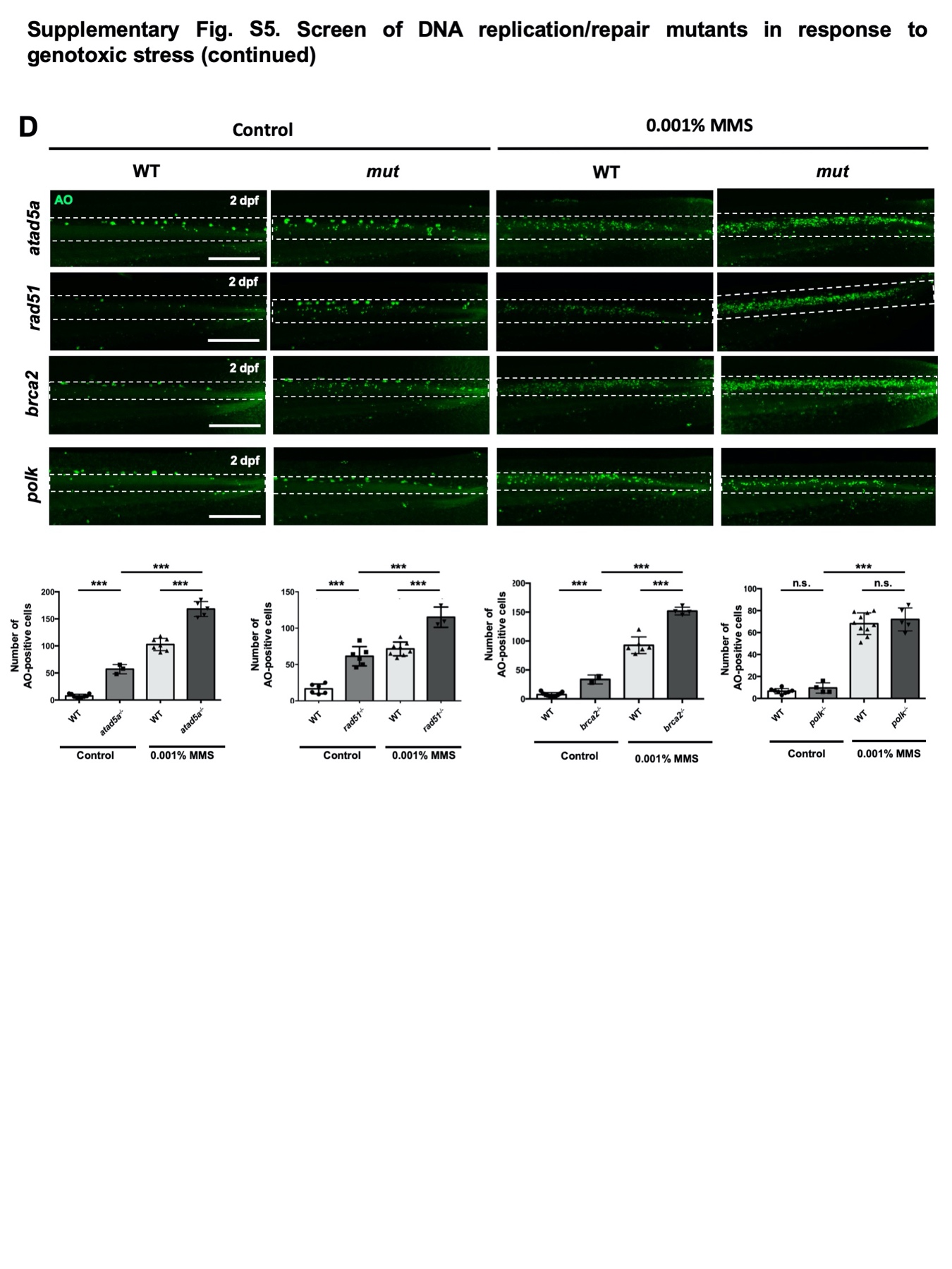

**Supplementary Fig. S5. Screen of DNA replication/repair mutants in response to genotoxic stresses. A.** Images of 4 or 5 dpf embryos generated from inbred heterozygous mutants of *rad51* and *shfm1* in response to 5Gy IR exposure at 6 hpf. Morphological alteration is indicated by red arrowheads in the homozygous mutants exposed with IR. **B.** Images of confocal microscopy with AO staining and quantification of AO–positive apoptotic cells in the brain of 4 dpf *rad51^-/-^* and in the ventral jaw structure of 5 dpf *shfm1^-/-^*(*cu46/cu46*) mutants with IR-induction. Yellow dotted area indicates dorsal view of the brain area and red dotted area indicates ventral view of the head structure. **C**. Images of 2 dpf progenies produced from inbred heterozygous mutant of *atad5a,* *rad51 brca2* and *polk* with treatment of 0.001% MMS from 6 hpf. **D**. Images of confocal microscopy showing AO-stained body trunk and their quantification of AO–positive apoptotic cells in the body trunk of 2dpf *atad5^-/-^*(*cu33/cu33*), *rad51^-/-^*(*cu54/cu54*), *brca2^-/-^*(*cu51/cu51*) and *polk^-/-^*(*cu13/cu13*), mutant embryos with their WT sibling controls in the presence of MMS. White dotted box indicates neural tissues in the trunk of embryos. All quantification graphs represent mean ± S.E.M. with individual values. p-values were calculated by unpaired two-tailed Student’s t-test. *** p<0.001, *p<0.05. n.s.; not significantly different. Black scale bar = 500 µm; white scale bar = 200 µm

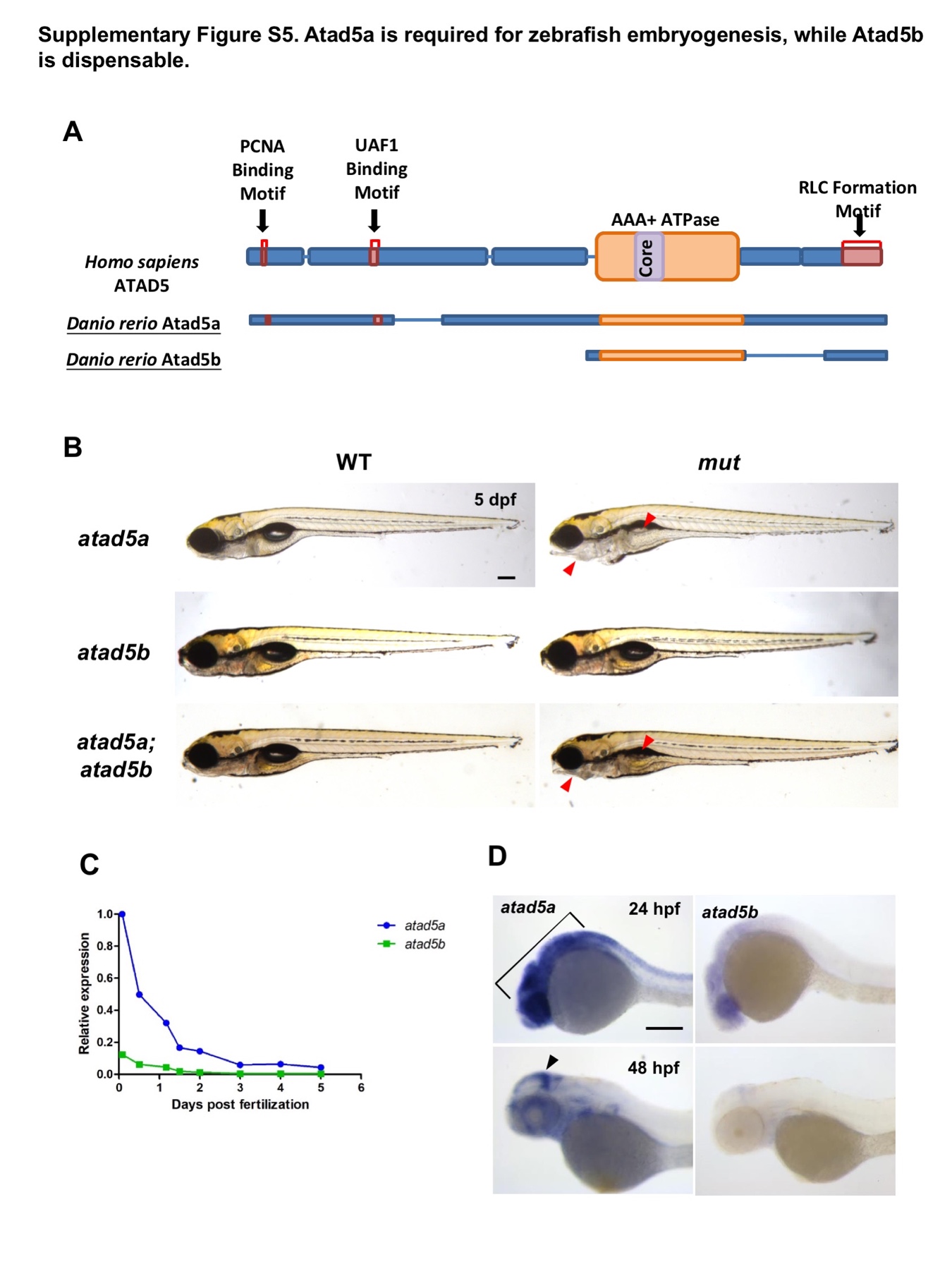
**Supplementary Fig. S6. Atad5a is required for zebrafish embryogenesis, while Atad5b is dispensable. A.** Diagrams describing functional domains of human ATAD5 protein and zebrafish paralogs, Atad5a and Atad5b. RLC stands for Replication factor C (RFC) Like Complex. **B.** Bright-field images displaying morphology of *atad5a^-/-^* single, *atad5b^-/-^* single and *atad5a^-/-^*; *atad5b^-/-^* double homozygous mutants at 5 dpf. Red arrowheads indicate abnormal appearance on the head and swim bladder in the mutant embryos. **C.** Quantitative real time polymerase chain reaction (qPCR) results of *atad5a* and *atad5b* during embryogenesis. **D**. Images of anterior part of zebrafish embryos with WISH using probes against *atad5a* and *atad5b* at 24 and 48 hpf. Black brackets and arrowhead indicate *atad5a*-positive tissues in the anterior part of the embryos. All results were normalized to 0 hpf of *atad5a*. *** p<0.001, **p<0.01, *p<0.05. Scale bar = 200 µm.

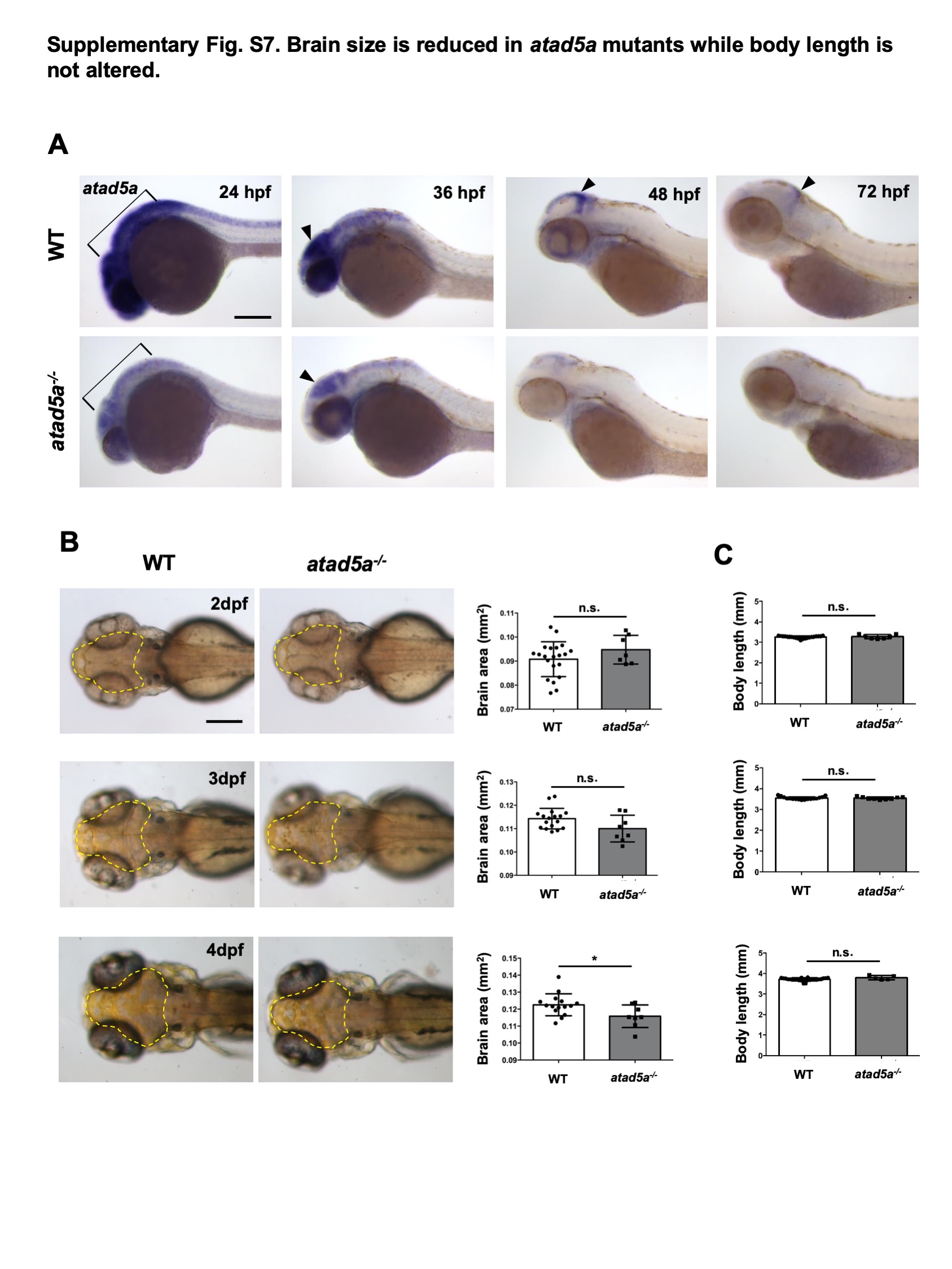

**Supplementary Fig. S7. Brain size is reduced in *atad5a* mutants while body length is not altered. A.** Expression of *atad5a* in the *atad5a^-/-^* and its WT sibling controls from 24 hpf to 72 hpf. Black brackets and arrowheads indicate *atad5a*-positive tissues in the anterior part of the embryos. **B.** Size assessment around forebrain to cerebellum (yellow dotted area) between *atad5a^-/-^* mutant and WT controls from 2 to 4 dpf and quantification of the brain size (yellow dotted area). **C.** Quantification of the entire body length between *atad5a^-/-^* mutant and WT controls from 2 to 4 dpf. Graphs represent mean ± S.E.M. with individual values. p-values were calculated by unpaired two-tailed Student’s t-test. *p<0.05. n.s.; not significantly different. Scale bar = 200 µm.

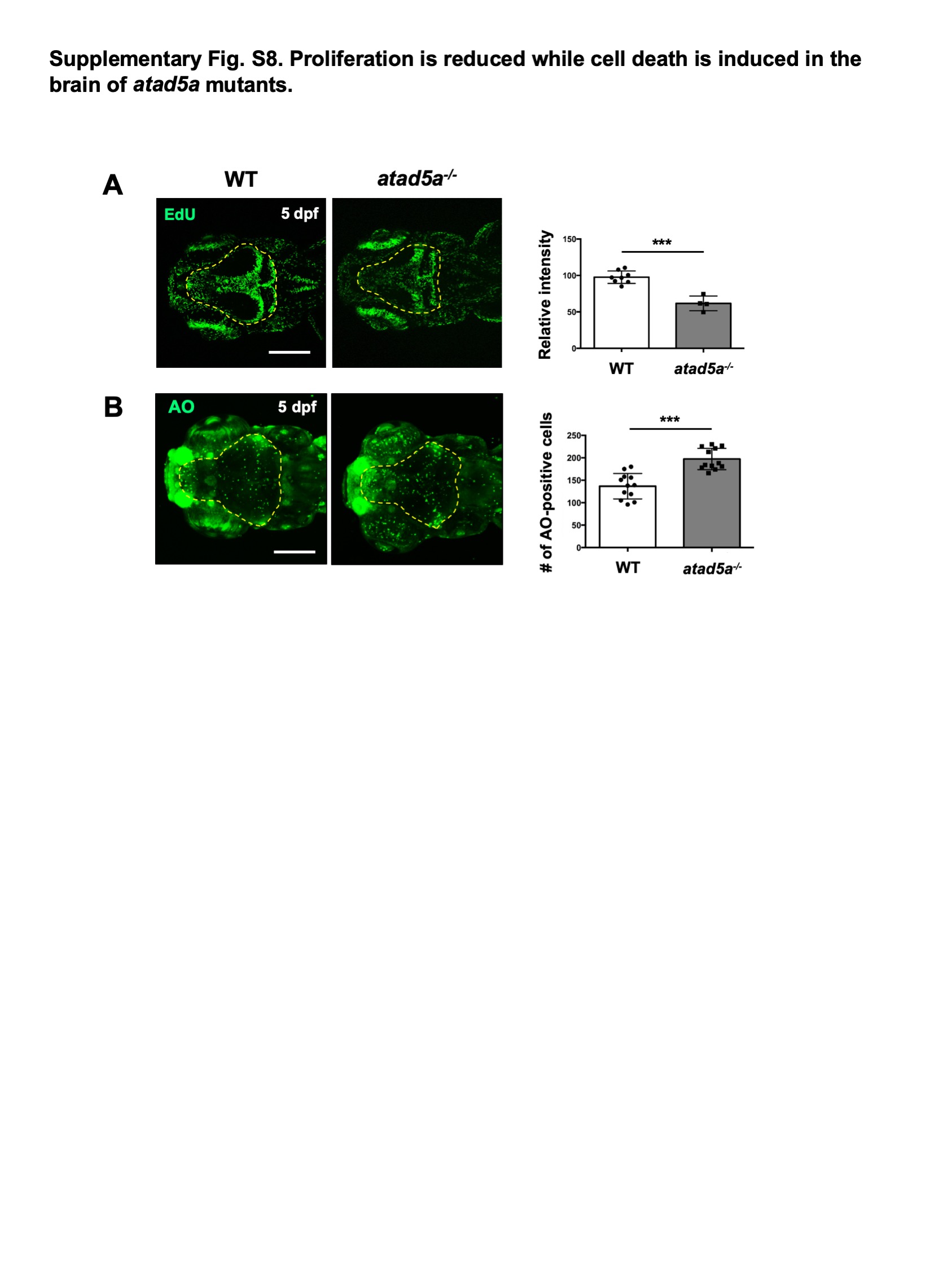

**Supplementary Fig. S8. Proliferation is reduced while cell death is induced in the brain of *atad5a* mutants. A.** Confocal microscope images of EdU-positive cells in the brain of *atad5a^-/-^* and WT controls at 5 dpf and quantification of relative intensity of EdU signal in the brain. Yellow-dotted area indicates the brain including forebrain to cerebellum. **B.** Confocal microscope images of AO-positive cells in the brain (dotted area) of *atad5a^-/-^* and WT controls at 5 dpf and quantification of AO-positive cell number in the brain. Graphs represent mean ± S.E.M. with individual values. p-values were calculated by unpaired two-tailed Student’s t-test. *** p<0.001. Scale bar = 200 µm.

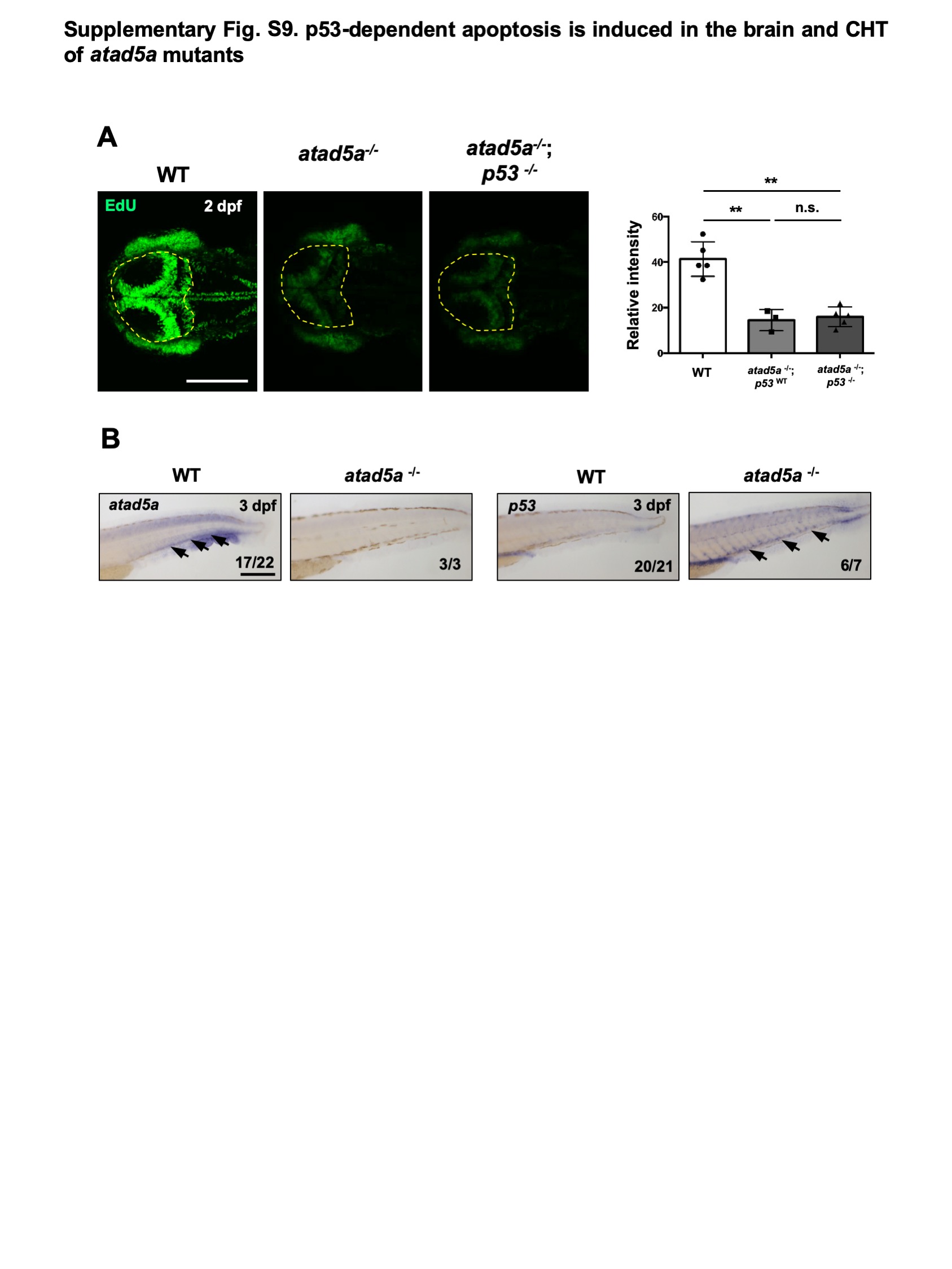

**Supplementary Fig. S9. p53-dependent apoptosis is induced in the brain and CHT of *atad5a^-/-^*** **mutants. A.** Confocal microscope images of EdU-positive cells in the brain (yellow dotted area) of *atad5a^-/-^* single, *atad5a^-/-^;p53^-/-^* double mutants and WT controls at 2 dpf and its quantification of relative intensity of EdU signal in the brain. Graphs represent mean ± S.E.M. with individual values. p-values were calculated by unpaired two-tailed Student’s t-test. **B.** Images of WISH with probes against *atad5a* and *p53* using *atad5a^-/-^* and WT controls at 3 dpf. Black arrows indicate violet WISH-positive signal in the CHT. **p<0.01, n.s.; not significantly different. Scale bar = 200 µm.

**Supplementary Table S1.** **Information on DNA repair genes targeted by CRISPR/Cas9 including injected sgRNA sequence, primers used for founder screening.**

*Supplementary Table S1 is included in Excel file.

```
1.apex1
wt			gaccagtaaggacggccgggcagccaatat (135-164)
```

*apex1^cu1^*(Δ2:-3+1)	gacca**gt--Tgacggccgggcagcc**aatat

*apex1^cu2^*(Δ4)	gacca**g----gacggccgggcagcc**aatat

```
2.unga
wt 			aaacagaagttggaaagcggaaatgaggca (100-129)
```

*unga^cu3^*(Δ5)	aaaca**gaagttggaaagcg-----g**aggca

```
3.ercc1
wt 			atccaaaaatgatcacgtagaggcctcatc (84-113)
```

*ercc1^cu4^*(Δ2)	atcca**--aatgatcacgtagaggcc**tcatc

*ercc1^cu5^*(Δ4)	atcca**aa----atcacgtagaggcc**tcatc

```
4.xpc
wt 			tcaaagaagaggaggaagacagtgaggatgaggacgact (296-334)
```

*xpc^cu6^*(Δ7)	tcaaagaaga**ggaggaagaca-------tg**aggacgact

*xpc^cu7^*(Δ8:-9+1)	tcaaagaaga**ggaggaagacagtgagg--------C**act

*xpc^cu8^*(Δ21:-24+3**)**	tca---------------------**TAAatg**aggacgact

```
5.neil1
wt			caaatgggagctggcaaaaaaacagaggccc (398-428)
```

*neil1^cu9^*(Δ14)	caaat**gggagct--------------**ggccc

*neil1^cu10^*(+8:-5+13)	caaat**gggagctggcaaaaaTGGCCCAAAAAAT**aggccc

```
6.polk
wt			ggtttggtgtccgtgctgccatgcccggattcattgcta (434-472)
```

*polk^cu11^*(Δ5)	ggttt**ggtgtccgtgctgcc-----**cggattcattgcta

*polk^cu12^*(Δ10:-25+15)	ggttt**ggtgt----------TAAAAGCTCTAAAAA**gcta

*polk^cu13^*(Δ13)	ggttt**ggtgtcc-------------**cggattcattgcta

```
7.rad18
wt 			tcaaaggcaaagcagcgagacagaagggcc (329-358)
```

*rad18^cu14^*(Δ4)	tcaaa**ggcaaagcagcgaga----a**gggcc

*rad18^cu15^*(+4:-2+6)	tcaaa**ggcaaagcagcgagacTTTTTTaa**gggcc

```
8.polh
wt			agggaggcgagtgtggaggtgatcgaggtg (276-305)
```

*polh^cu16^*(Δ9)	aggga**ggcgagtgtgg---------**aggtg

```
9.shprh
wt			cgcttggagtctggcctcatcgaggaggga (823-852)
```

*shprh^cu17^*(+1):	cgctt**ggagtctggcctcatcgaAgg**aggga

```
10.hltf
wt			tcggagggattttggctgatgatatggggc (821-850)
```

*hltf ^cu18^*(Δ2)	tcgga**gggattttggctgatgat--**ggggc

```
11.msh2
wt 			tgtcggtgtggtgggtgttcgtcttggcactggcactgat (429-468)
```

*msh2^cu19^*(Δ16)	tgtc**ggtgtggtgggt----------------**gcactgat

*msh2^cu20^*(Δ3)	tgtc**ggtgtggtgggtgttc---t**tggcactggcactgat

*msh2^cu21^*(Δ30)	tgtc**gg------------------------------**tgat

```
12.mutyh
Wt 			ggtcaggtcttggatactactccagaggacgta (410-442)
```

*mutyh^cu22^*(Δ1)	ggtca**ggtcttggatactactc-ag**aggacgta

*mutyh^cu23^*(Δ10)	ggtca**ggtcttggatact----------**acgta

```
13.fanci
wt 			ttcagggatgttcccttatcaacggaggagc (556-586)
```

*fanci^cu24^*(Δ7:-22+15)	ttcag**g**-------**AGGAGGAGGAGCTTC**agc

*fanci^cu25^*(Δ10)	ttcag**ggatgttc----------gg**aggagc

```
14.usp1
wt 			ttcctctgtggagcagctgcagtccagctt (468-497)
```

*usp1^cu26^*(Δ2:-4+2)	ttcct**ctg--CTgcagctgcagtcc**agctt

```
15.palb2
wt 			aaagaggcagaggaagaggacggagagggaggagagg (1184-1220)
```

*palb2^cu27^*(Δ18)	aaaga**ggcagagga------------------**agagg

```
16.pcna
wt			agtatggactcctctcatgtgtctctggtgca (115-146)
```

*pcna^cu28^*(Δ1)	agtat**ggactcctctcatgtg-ctc**tggtgca

*pcna^cu29^*(Δ7)	agtat**ggactcctctcatgt-----**--gtgca

*pcna^cu30^*(Δ6)	agtat**ggactcctctcatg------**tggtgca

*pcna^cu31^*(Δ9)	agtat**ggactcctctc---------**tggtgca

*pcna^cu32^*(+9)	agtat**ggactcctctcatgtgtGGATGTCCTctc**tggtgca

```
17.atad5a
wt			ttaaaggagttctgtaatataccatcggtg (2380-2409)
```

*atad5a^cu33^*(Δ1)	ttaaa**ggagttctgtaatatac-at**cggtg

```
18.atad5b
wt			gccctgcgctgtggtcgactcctcctttct (1435-1464)
```

*atad5b^cu34^*(Δ2)	gccct**--gctgtggtcgactcctcc**tttct

*atad5b^cu35^*(Δ7)	gccct-------**ggtcgactcctcc**tttct

```
19.ino80
wt 			cagacggagctgtacgctcactttatgggt (1141-1170)
```

*ino80 ^cu36^*(Δ2)	cagac**ggagctgtacgctca--tta**tgggt

```
20. atrip
wt			catcaggagcgcgagcagcagcggcaggcg (457-486)
```

*atrip^cu37^* (+7:-3+10)	catca**ggagcgcgagcagcagTTTAAATACCc**aggcg

*atrip^cu38^* (Δ3)	catca**ggagcgcgagcagcagc---**aggcg

*atrip^cu39^* (Δ6)	catca**ggagcgcgagcagc------**aggcg

```
21. telo2
wt 			tcacaggactcatacgcagcctcagtgg (788-815)
```

*telo2^cu40^* (Δ2)	tcaca**ggactcatacgcagcc--ag**tgg

*telo2^cu41^* (Δ8)	tcaca**ggactcatacgcag--------**g

```
22. rtel1
wt			gaaagggaagacaccatggttggggaggaa (2560-2589)
```

rtel1*^cu42^* (Δ7)	gaaag**ggaagacaccatg-------**aggaa

```
23. ddb1
wt			ctcaaggccttcaacatccggctggaggag (454-483)
```

*ddb1^cu43^* (+2)	ctcaa**ggccttcaacatccggcCTtgg**aggag

```
24. hmgb1a
wt			aaactggacaaggcacgttacgagagggag (190-219)
```

*hmgb1a^cu44^*(Δ1:-4+3)	aaact**ggacaaggcacgtGTT-aga**gggag

*hmgb1a^cu45^*(Δ2)	aaact**ggacaaggcacgttacga--**gggag

```
25. shfm1
wt			cttttggaggaggatgatgaatttgaggaa (100-129)
```

*shfm1^cu46^* (Δ8)	ctttt**ggaggaggat--------tg**aggaa

```
26. nudt1
wt			gagctggaaaatggaacggctttggaggaa (83-112)
```

*nudt1^cu47^* (Δ2)	gagct**ggaaaatggaacggct--gg**aggaa

*nudt1^cu48^* (+5, Δ1+6)	gagct**ggaaaatggaacggctGGAAAAtgg**aggaa

*nudt1^cu49^* (Δ6)	gagct**ggaaaatggaacgg------**aggaa

```
27. brca2
wt			tctttggtagacaataacaacacggtggca (1051-1080)
```

*brca2^cu50^*(Δ5)	tcttt**ggtagacaataac-----gg**tggca

*brca2^cu51^*(+2:-1+3)	tcttt**ggtagacaataacaacacCCAg**tggca

*brca2^cu52^*(+21:1-+22)	tcttt**ggtagacaataacaacaTGTGGCACCCAGTGAAACTAATgg**tggca

```
28. blm
wt 			aaccgaagacaaatacagttaatccactag (293-322)
```

*blm^cu53^* (+5:-1+6)	aaccg**CTATTTagacaaatacagttaatcc**actag

```
29. rad51
wt			caccagcgcagagccgaaatcatccagatc (280-309)
```

*rad51^cu54^*(Δ7)	cacca**-------gccgaaatcatcc**agatc

*rad51^cu55^*(Δ3)	cacca**gc---gagccgaaatcatcc**agatc

```
30. mre11a
wt			atggaggtgatgagaaagtactgcatgggg (229-258)
```

*mre11a^cu56^*(+1)	atgga**ggtgatgagaaagtactTgca**tgggg

```
31. xrcc1
wt 			gtgaaggcgcaatcacaggagaagtgggatcgcg (346-379)
```

*xrcc1^cu57^*(Δ1, Δ3+2)	gtgaa**ggcgcaatcacagga-TTgt**gggatcgcg

*xrcc1^cu58^*(Δ7)	gtgaa**ggcgcaatcacaggaga-------**tcgcg

*xrcc1^cu59^*(+1:-3+4)	gtgaa**ggcgcaatcacaggagaTGCC**gggatcgcg

*xrcc1^cu60^*(+11:-4+15)	gtgaa**ggcgcaatcacagTGAAGGCGCAATCACagt**gggatcgcg

*xrcc1^cu61^*(Δ3)	gtgaa**ggcgcaatcacaggag---t**gggatcgcg

```
32. rad54l
wt 			cttgtgggattgtgtgacgggacgacggat (471-500)
```

*rad54l^cu62^*(Δ8)	cttgt**gggattgtgtga--------**cggat

*rad54l^cu63^*(Δ3)	cttgt**gggattgtgtgacggga---**cggat

```
33. wrn
wt			gaacaggccagacaggagcagctggaggag (1036-1065)
```

*wrn^cu64^*(+2:-2+4)	gaaca**ggccagacaggagcagTGTAgg**aggag

*wrn^cu65^*(Δ9)	gaaca**ggccagacagg---------**aggag

```
34. slx1b
wt 			accgaggaggagccaagagaaccagtggca (134-163)
```

*slx1b^cu66^*(Δ1)	accga**ggaggagccaagagaac-ag**tggca

*slx1b^cu67^*(+5:-2+7)	accga**ggaggagccaagagaacGTGGAAGg**tggca

**Supplementary Table S2. List of mutant alleles of DNA repair genes and each sgRNA target sequence.** The number in parentheses with the wild-type (wt) sequences indicates the position in the open reading frame of the target sequences in each gene. The sgRNA sequence is shown in bold letter. Deletions are illustrated as dashed lines and insertions as capital letters in blue.

**Supplementary Table S3.** **Allele information for genotyped animals for mutant phenotyping**

*Supplementary Table S3 is included in Excel file.

**Supplementary Table S4.** **Fertility test for the mutants with defects of sex reversal.**

*Supplementary Table S4 is included in Excel file.

**Supplementary Table S5.** **Genotyping result of screening for the sensitivity against genotoxic stress.**

*Supplementary Table S5 is included in Excel file.
